# A real-time, high-performance brain-computer interface for finger decoding and quadcopter control

**DOI:** 10.1101/2024.02.06.578107

**Authors:** Matthew S. Willsey, Nishal P. Shah, Donald T. Avansino, Nick V. Hahn, Ryan M. Jamiolkowski, Foram B. Kamdar, Leigh R. Hochberg, Francis R. Willett, Jaimie M. Henderson

## Abstract

People with paralysis express unmet needs for peer support, leisure activities, and sporting activities. Many within the general population rely on social media and massively multiplayer video games to address these needs. We developed a high-performance finger brain-computer-interface system allowing continuous control of 3 independent finger groups with 2D thumb movements. The system was tested in a human research participant over sequential trials requiring fingers to reach and hold on targets, with an average acquisition rate of 76 targets/minute and completion time of 1.58 ± 0.06 seconds. Performance compared favorably to previous animal studies, despite a 2-fold increase in the decoded degrees-of-freedom (DOF). Finger positions were then used for 4-DOF velocity control of a virtual quadcopter, demonstrating functionality over both fixed and random obstacle courses. This approach shows promise for controlling multiple-DOF end-effectors, such as robotic fingers or digital interfaces for work, entertainment, and socialization.

---

More than 5 million people in the United States live with severe motor impairments^1^. Although many basic needs of people with paralysis are being met, unmet needs for peer support, leisure activities, and sporting activities are reported, respectively, by 79%, 50%, and 63% of surveyed people with paralysis from spinal cord injury.^2^ People with motor impairments that spare enough function to manipulate a video game controller have turned to video games for social connectedness and a competitive outlet^3,4^. In a survey of players with and without disabilities^3^, a variety of themes emerged (e.g., recreation, artistic expression, social connectedness); however, in those with disabilities – in contrast to those without – many expressed a theme of enablement, meaning both equality with able-bodied players and overcoming their disability. Even with assistive/adaptive technologies, gamers with motor impairments often have to play at an easier level of difficulty^5^ or avoid multiplayer games with able-bodied players^6^ that often require dexterous multi-effector control.^4,7^ Brain-computer interfaces (BCIs) could enable sophisticated control of video games for people with paralysis – and more broadly, control of digital interfaces for social networking or remote work. BCIs are being increasingly recognized as a potential solution for motor restoration and have been used for controlling a robotic arm with somatosensory feedback^8^; controlling computer tablets^9^ and cursors^10^; decoding handwriting^11^; producing text^12^; and synthesizing speech^13^.

In motor BCIs, most effort has focused on controlling single effectors such as computer cursors for point-and-click cursor control and robotic arms for reaching/grasping (where fingers moved as a group)^10,14-17^. However, fine motor restoration with dexterous finger control would allow better object manipulation, and could enable activities such as typing, playing a musical instrument, or manipulating a multi-effector digital interface such as a video game controller. To expand object manipulation, Wodlinger et al.^18^ continuously decoded linear combinations of 4 distinct grasping postures, although fingers themselves were not individuated. Furthermore, several groups have shown that finger movements can be differentially classified using neural activity recorded from BCIs in human research participants^19-21^. Recent work in non-human primates (NHPs) has moved beyond simple classification, demonstrating continuous decoding for the dexterous (i.e., individuated) control of two individuated finger groups^22,23^. Expanding on this work, we developed the most capable finger BCI system to date adapting recently introduced non-linear decoding techniques^23^, providing continuous decoding of three finger groups with 2D thumb movements (4 total degrees of freedom (DOF)) in a human research participant with paralysis – doubling the decoded degrees of freedom in animal studies^22,23^. We determined how the dimensionality of the neural representation increases with the number of decoded fingers and also estimated the dependence of decoding accuracy on the number of recording electrodes. Finally, we used the decoded finger movements to provide independent digital endpoints that control a virtual quadcopter with 4 DOF to demonstrate that intracortical BCIs (iBCIs) can allow a multi-effector, high-throughput brain-to-digital connection for video game play. This system allows an intuitive framework to control a digital interface (similar to a video game controller) that can be broadly applied to a variety of control applications including remote work and recreation.

## Results

Multi-unit neural activity was recorded from two 96-channel silicon microelectrode arrays placed in the hand ‘knob’ area of the left precentral gyrus in one participant (‘T5’) enrolled in the BrainGate2 pilot clinical trial (Extended Data Fig. 1a). A virtual hand was displayed to the participant using Unity (version 2021.3.9f1, Unity Technologies, San Francisco, CA, USA), as shown in Fig. 1a. The thumb was designed to move along a 2-dimensional surface defined by the flexion-extension and abduction-adduction axes (Fig. 1b). Both the index-middle and ring-small fingers moved as separate groups in a 1-dimensional arc constrained to the flexion-extension axis.

**Figure 1.**
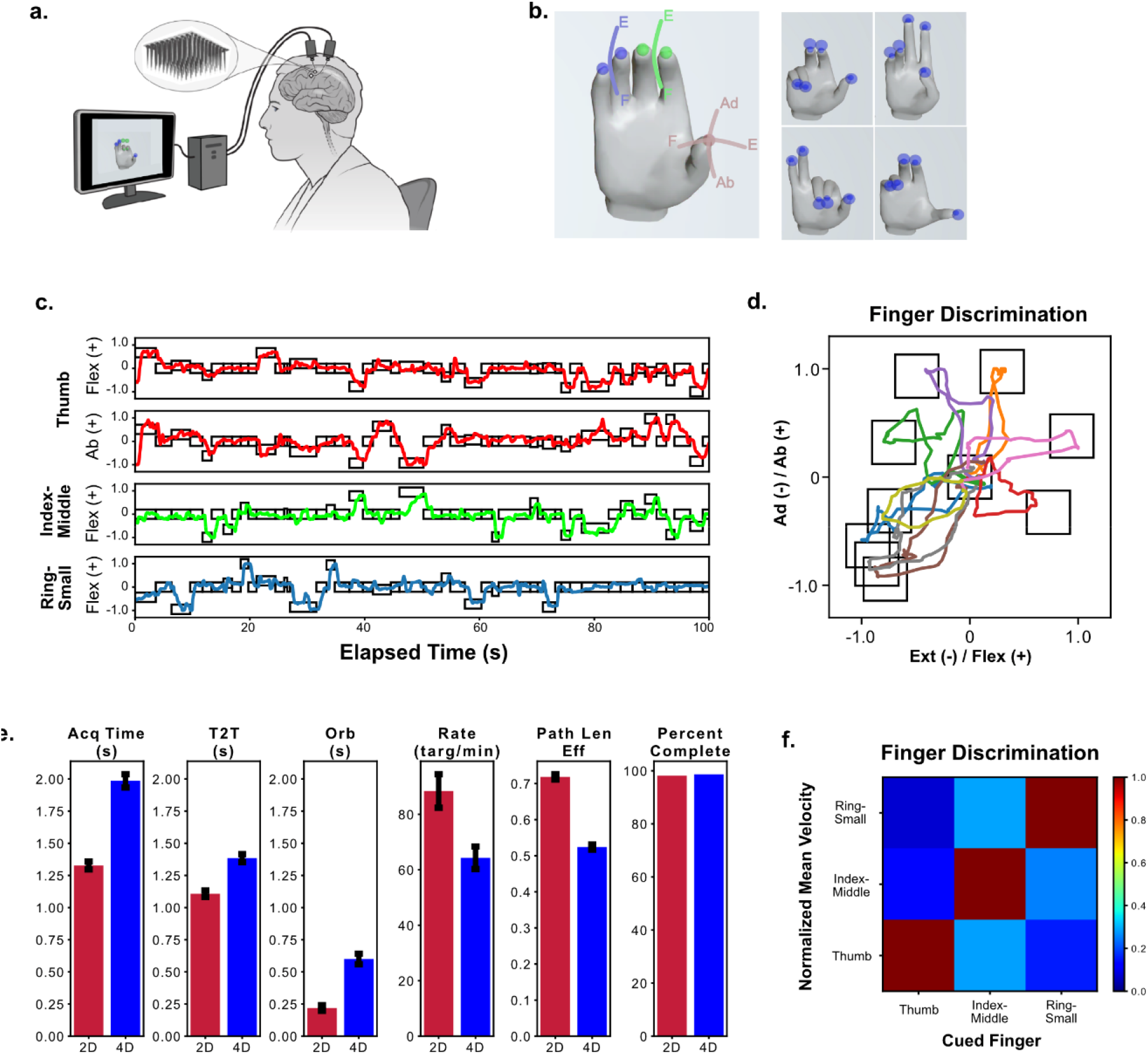
Intracortical brain-computer interface system for dexterous finger movements. **a**, A computer display is placed in front of the participant so that he can perform a finger task with a virtual hand. During closed-loop control, the electrical activity from the array is mapped to a control signal for the virtual fingers. Panel adapted from Willet et al. (2021)^11^. **b (left)**, Thumb moves in two dimensions, abduction (Ab) and adduction (Ad) (flexion/extension and abduction/adduction), while index-middle and ring-small move in a 1D arc. F, flexion; E, extension. **b (right)**, Trials showing typical targets for all 3 finger groups for the 4-degree-of-freedom task. **c**, A 100-s time segment of typical decoded movements are depicted for the 3-finger group, 4-degree-of-freedom (DOF) task. Trajectories are described along a range of -1 to 1, where 1 denotes full flexion or abduction and -1 denotes extension or adduction. **d**, The 2D thumb trajectories for the full 50-target block shown in **c**. Titles for each panel indicate the 2D thumb distance from the neutral position in arbitrary units (where 1 is 100% 1D flexion/abduction and 0 is 100% 1D extension/adduction). **e**, Summary statistics comparing the 2- and 4-DOF tasks for acquisition time (Acq Time), time to target (T2T), orbiting time (Orb), acquisition rate (Rate), path length efficiency (Path Len Eff), and the percent of trials successfully completed (Percent Complete). The error bars represent the standard error of the mean. **f**, Four blocks in which only 1 finger was cued per trial, illustrating individuated control of fingers. The mean velocity per trial was calculated during the ‘Go’ period for each finger and normalized by the mean value of the finger group with the highest mean value.

### Closed-loop, real-time control of a 2- and 4-DOF finger task

To perform closed-loop continuous decoding, a temporal-integrated feed-forward neural network, adapted from Willsey et al.^23^, mapped spike-band power (SBP)^24^ to finger velocities used to control virtual finger movements on screen (Extended Data Fig. 2). Two sets of tasks were performed. First, we sought to translate findings from earlier NHP studies^22,23^ demonstrating decoding of two finger groups (2D task) to our human research participant (where in this task the thumb was constrained only to the flexion-extension axis). T5 was cued to move both the thumb and index-middle groups from a center position to a random target within the active range of motion of the fingers. On the subsequent trial, the targets were placed back at the center. To successfully complete a trial, the fingers had to hold on the targets for 500 ms, and 10 s were allowed to complete the trial (sample trajectories in Extended Data Fig. 3a; see Movie 1).

To expand on the functionality demonstrated in NHP studies, task complexity was increased by introducing a 4D task with 2D thumb movements and 1D movements of the index-middle group and the ring-small group (Fig. 1c). On each trial, 2 finger groups were randomly selected for new targets while the target for the third finger group remained in the same position as the previous trial, and movements of all fingers were continuously and simultaneously decoded and controlled. Typical target trajectories for this expanded 4D task are shown in Fig. 1c, and 2D trajectories of the thumb movements are illustrated in Fig. 1d. A video of this task is shown in Movie 2.

The closed-loop decoding performance for the 2D and 4D decoder was compared using 529 trials (3 days) for the 2D decoder and 524 trials (6 days) for the 4D decoder (Fig. 1e). For the 2D decoder, the mean acquisition time was 1.33 ± 0.03 s, the target acquisition rate was 88 ± 6 targets/min, and 98.1% of trials were successfully completed. For the 4D decoder, the mean acquisition time was 1.98 ± 0.05 s, target acquisition rate was 64 ± 4 targets/min, and 98.7% of trials were successfully completed. The acquisition times for each trial (population data) is shown graphically in Extended Data Fig. 3b for the 2D decoder and Extended Data Fig. 3c for the 4D decoder. Typical finger distances per trial are shown graphically in Extended Data Fig. 4a for the 2D task and Extended Data Fig. 4b for the 4D task.

In comparison to the 2D decoder and task, the acquisition times were increased by 50% for the 4D decoder and task (*P* < 10^−10^, *t* = -11.00, *df* = 1051, *CI* = -774 to -540). However, after the participant grew more accustomed to the task (final 4 blocks), acquisition time for the 4D decoder dropped by an average of 0.4 s to 1.58 ± 0.06 s (a target acquisition rate of 76 ± 2 targets/min), and 100% of trials were completed. To compare this work with the previous NHP 2-finger task where throughput varied from 1.98 to 3.04 bps with a variety of decoding algorithms^23,25^, throughput for the current method was calculated as 2.64 ± 0.09 bps (see Methods for details). Table 1 summarizes statistics for the 4D decoder/task and 2D decoder/task.

**Table 1:** Performance metrics for 2D and 4D finger decoding.

|  | 2D Decoder | 4D Decoder | 4D Decoder<br>(last 4 runs) |
| --- | --- | --- | --- |
| Number of trials | 529 | 524 | 192 |
| Number of days | 3 | 6 | 3 |
| Acquisition time (ms) | 1330 ± 30 | 1980 ± 50 | 1580 ± 60 |
| Time to target (ms) | 1110 ± 30 | 1380 ± 30 | 1200 ± 40 |
| Orbiting time (ms) | 220 ± 20 | 600 ± 40 | 380 ± 50 |
| Targets per minute | 88 ± 6 | 64 ± 4 | 76 ± 2 |
| Percent completed | 98.1% | 98.7% | 100% |
| Path length | 0.718 ± 0.007 | 0.524 ± 0.007 | 0.580 ± 0.010 |

Statistical data are reported as mean ± standard error of the mean.

To graphically illustrate the degree of finger discrimination during closed-loop control, the 4D decoder was also run on a 4D task in which only 1 finger group was cued to move on each trial (1 finger group had a new target and the other 2 finger groups had the same target as the previous trial). During movement of the cued finger group, the mean velocity of the non-cued finger groups was calculated during the ‘Go’ period of the trial. The movement of the non-cued fingers was substantially less than the movement of the cued finger (Fig. 1f), demonstrating that finger groups can be individuated to allow for dexterous finger tasks.

To understand how simplifying the task complexity would affect the trade-off between acquisition time and target acquisition rate, the 4D decoder was compared for tasks with 1 and 2 cued finger-group movements on each trial. For 1 cued finger (178 trials), the mean acquisition time was 1.37 ± 0.06 s and target acquisition rate was 45 ± 4 targets/min (see Movie 3), and for 2 cued fingers (187 trials), the mean acquisition time was 1.66 ± 0.07 s and target acquisition rate was 74 ± 6 targets/min (see Extended Data Fig. 5a and Extended Data Table 1). Thus, cueing 1 finger group led to shorter trial times (*P* = 0.0036, *t* = -2.93, *df* = 363, *CI* = -485.4831 to -95.7223), and cueing 2 finger groups led to higher target acquisition rate (*P* = 0.0092, *t* = -3.78, *df* = 6, *CI* = -47.7206 to -10.1928).

### Dimensionality of the neural activity

Intuitively, one would expect that as the dimensionality of decoded DOF increases, the dimensionality of the neural activity should also increase. To determine the relationship between the dimensionality of the neural activity and decoded DOF, the dimensionality of the neural data during 4D and 2D decoding was calculated using the metric defined by Willett et al.^11^, which uses the participation ratio to quantify dimensionality (see Methods). The average dimensionality of neural activity was 2.4 for the 2D decoder, 3.1 for the 4D decoder with 1 new target/trial, and 7.5 for the 4D decoder with 2 new targets/trial (Fig. 2a). If the dimensionality of the neural activity varied linearly with the decoded DOF, the dimensionality of the 4D decoder would be twice that of the 2D decoder, i.e., 2 × 2.4 = 4.8; however, dimensionality using the 4D decoder was found to be 7.5, 56% more than the expected value of 4.8 (*P* = 0.028, *t* = -2.77, *df* = 7, *CI* = -4.9901 to -0.3908). Thus, the dimensionality of combined finger movements was greater than the sum of the individual components, implying that some neurons may encode both single movements and movement combinations.

**Figure 2.**
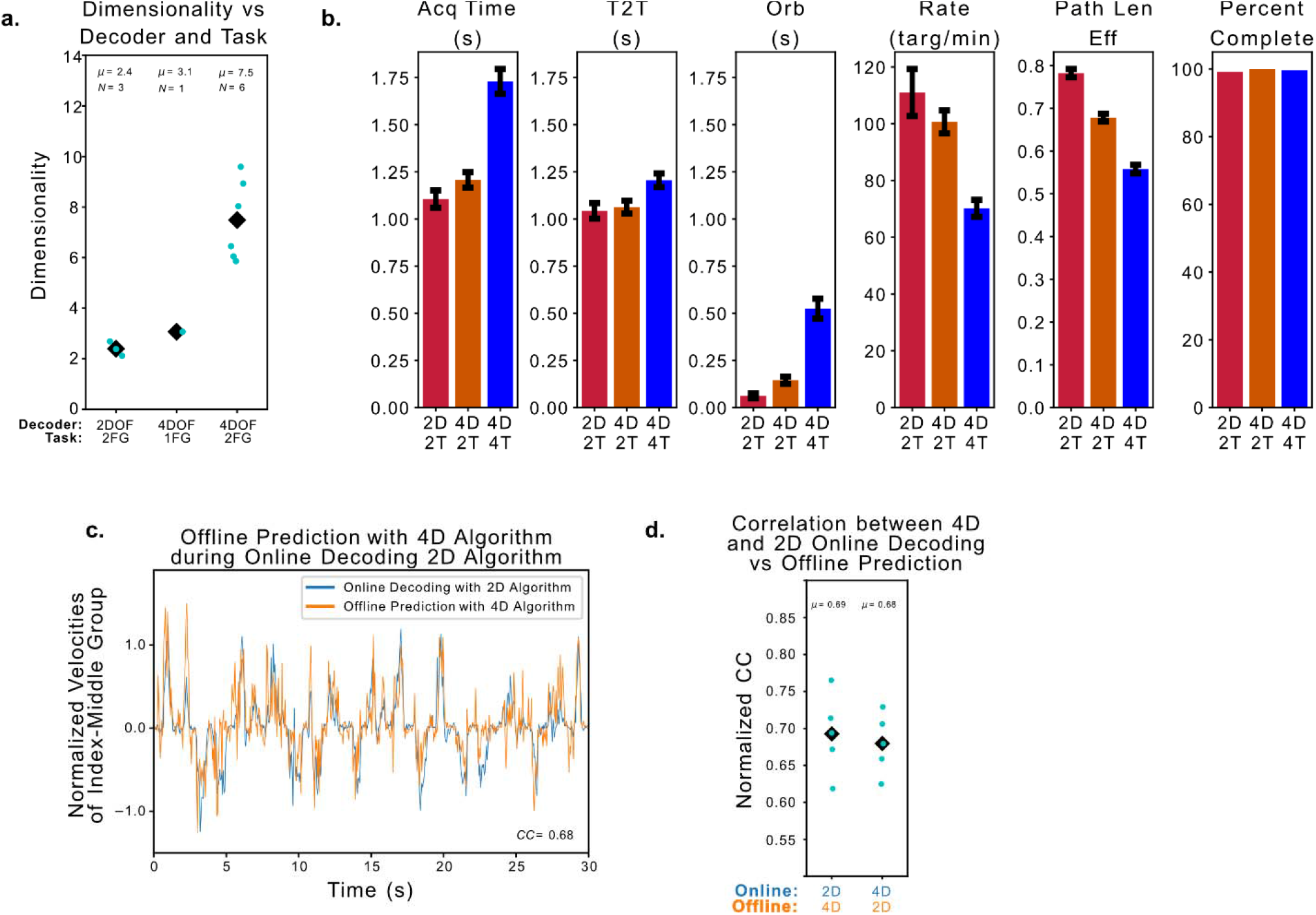
Dimensionality and dimensionality reduction. **a**, The dimensionality of the neural activity during closed-loop decoding using either the 2D decoder and task or 4D decoder and task for either 1 or 2 new finger-group targets (1 FG or 2 FG) per trial. Each light blue dot represents data from a single day, and the black diamond represents the mean value. The mean, μ, is given for each decoder/task pairing. DOF, degree of freedom. **b**, Summary statistics comparing the 2D decoder on the 2D task (2D, 2T), the 4D decoder on the 2D task (4D, 2T), and the 4D decoder on the 4T task (4D, 4T) based on the acquisition time (Acq Time), time to target (T2T), orbiting time (Orb), acquisition rate (Rate), path length efficiency (Path Len Eff), and the percent of trials successfully completed (Percent Complete). The error bars represent the standard error of the mean. **c**, A typical online block showing the decoded index-middle finger group velocities using the 2D on 2T (blue) during an online block. Offline, the 4D decoding algorithm was used to predict index-middle group velocities from the same block (orange). The normalized cross correlation (CC) between the online and offline signals, which quantifies the similarities between the signals, is given in the bottom right corner. The units of velocity are calculated using a distance of 1 to denote the full active range of motion. **d**, For the 10 blocks on the 2D task, the 4D decoding algorithm was used to predict finger velocities during online blocks using the 2D decoder (online 2D in blue, offline 4D in orange), and the 2D decoding algorithm was used to predict online velocities using the 4D decoder (online 4D in blue, offline 2D in orange). Each dot represents the normalized cross-correlation value between the offline and online signals averaged across both finger groups. The average of all 5 blocks for each paired comparison is indicated with a black diamond, and μ denotes the mean value.

### Effect of number of active DOF on decoding

Even though there is increased dimensionality in neural activity when decoding more DOF, it is unclear whether decoding more DOF impacts the mapping of the neural activity when decoding a lower number of DOF. The neural representation of the DOF decoded in the 2D task (thumb and index-middle flexion/extension) could change during the 4D task, for example, if a different control strategy is required for the 4D compared to the 2D task – similar to how new control strategies can be developed to account for a perturbation in the mapping from neural activity to the DOF^26^. Alternatively, the original neural representation could be suppressed when tasked with decoding additional fingers, as is the case when decoding unilateral vs bilateral movements^27^. A third competing hypothesis is that the neural representation of finger movement in the 2D task is preserved in the 4D task, similar to preservation of neural representation between open-loop motor imagery and closed-loop control^28^.

To explore these hypotheses, 2D and 4D decoders were trained and compared by testing on the 2 shared DOF (thumb flexion/extension and index-middle group flexion/extension) over 2 days (662 trials), in alternating trials (see Fig. 2b, Extended Data Table 2). The mean acquisition time was 1.11 ± 0.05 s for the 2D decoder on the 2D task (N = 233), 1.73 ± 0.07 s for the 4D decoder on the 4D task (N = 284), and 1.21 ± 0.04 s for the 4D decoder on the 2D task (N = 329). A movie of the 4D decoder on the 2D task is shown in Movie 4. The trial-by-trial acquisition times for this comparison are given in Extended Data Fig. 5c. The 4D decoder performed much closer to the 2D decoder when restricted to the same 2D task (9.2% increased acquisition times, *P* = 0.10, *t* = -1.64, *df* = 560, *CI* = -224.3189 to 20.1014). Thus, training a decoder on an expanded set of movements does not appear to substantially degrade decoding performance (summarized in Fig. 2b and Extended Data Table 2).

The mapping from neural activity to the original 2 DOF was compared for both 2D and 4D decoders. To do this, the 4D decoder was used to predict the velocities decoded by the 2D decoder on the 2D task and vice versa. The predicted velocities from the 4D decoder were similar to those decoded in online blocks by the 2D decoder (Fig. 2c). To quantify this comparison, the normalized cross-correlation (CC) function was calculated between the decoded and predicted velocities during the 8 blocks (Fig. 2d). The results were separated based on whether the online decoded velocities were from the 4D or 2D decoders. The CC when the 4D algorithm predicted the 2D decoded velocities was 0.69 ± 0.02, and when the 2D algorithm predicted the 4D decoded velocities, the CC was 0.68 ± 0.02 (Fig. 2c). Thus, both the 2D and 4D decoding algorithms had similar mapping when reducing the dimensionality of the original input channels.

### Dependency of decoding accuracy on channel count

Newer implantable BCI devices are expected to have more input channels than the device used in this study. To explore whether increasing the channel count could increase decoding accuracy, a vector-based, sample-by-sample signal-to-noise ratio (SNR) metric was formulated (Fig. 3A). In this formulation, the predicted/decoded finger velocities are compared with idealized velocities inferred from intended finger movements. The component of the predicted/decoded velocities consistent with idealized velocities (i.e., the component parallel to the idealized vector of finger velocities) is considered the signal component, while the predicted/decoded velocities inconsistent (i.e., the component orthogonal to the idealized velocity vector) are considered noise. The ratio of the expected signal mean over the square root of the noise power was denoted as the directional SNR (dSNR).

**Figure 3.**
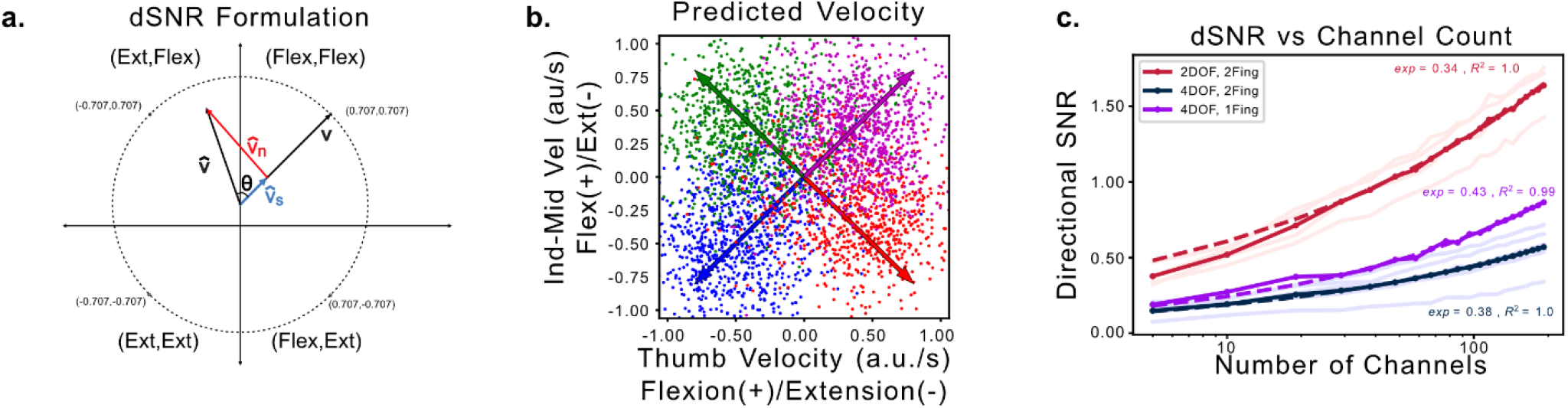
Signal-to-noise ratio vs. channel count. **a**, Graphical representation for vectorized directional signal-to-noise ratio (dSNR) for the 2D decoder and task. The positive *x* axis represents velocities flexing the thumb and negative values represent velocities extending the thumb. The *y* axis represents velocities flexing the index-middle finger group when positive and extending the finger group when negative. Since the signal vector, ***v***, is assumed to be a normalized target vector, the values of ***v*** can be only the 4 points indicated on the circle. The decoded/predicted velocity, 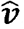, will lie at an angle θ to ***v*** and can be decomposed into a parallel signal component, 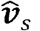, and a perpendicular noise component, 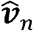. These components can be used to calculate dSNR. **b**, Velocities predicted by linear regression (using all *Nc* = 192 channels) that maps neural activity to finger velocities, which together with the intended finger movements are used to calculate dSNR. The arrows represent the ideal/truth value for each possible finger position based on the assumed intended finger movement. au, arbitrary unit. **c**, The dSNR as a function of channel count for the 2D decoder on the 2-target/trial task (red), 4D decoder on the 2-target/trial task (blue), and 4D decoder on the 1 target/trial task (purple). An empirical calculation of dSNR for each day is depicted in lightly colored lines and the mean value as the dark solid line. The dashed lines correspond to a linear, least-squares fit for the log-log relationship in Eq. 1, where *m* denotes the log-log slope and *R*^*2*^ denotes the coefficient of determination of the linear fit.

The value of dSNR was calculated during a “go” period of closed-loop trials (defined as 200-700 ms from trial onset) of 2- and 3-finger decoding (Extended Data Table 4). On each day, linear regression was used to train a mapping (against the intended finger direction) to convert SBP to finger velocities (using 6-fold cross-validation). Predicted velocities (calculated from all 192 input channels) grouped along the idealized, intended directions (Fig 3b).

To determine the dependency dSNR on channel count, a linear mapping of SBP to velocities was trained for a given number of *N* channels, which was used to calculate dSNR (using 6-fold cross-validation and where dSNR was the average using 25 sets of *N* randomly-selected channels; see Methods for details). For both the 2D and 4D tasks requiring movement of 2 simultaneous finger groups, dSNR did not saturate with increasing numbers of input channels (Fig. 3c). Since the dSNR metric assumes that both finger groups are simultaneously moving toward their respective targets (as opposed to moving one at a time), a simpler 4D task that required only 1 cued finger movement/trial was also performed (Fig. 3c). Using the dSNR data for the highest 75% of channel counts of each curve, a log-log relationship between channel count and dSNR was empirically fit to a linear relationship. The empirical fit of the log-log relationship was strongly linear, with a coefficient of determination, *R*^*2*^, between 0.99-1.00 and a slope, *m*, of 0.34 for the 2D task moving 2 fingers, 0.38 for the 4D task moving 2 fingers, and 0.43 for the behaviorally simpler 4D task moving 1 finger. Given the high *R*^*2*^ value, the empirical relationship between the dSNR and channel count fit the relationship in Eq. 1,

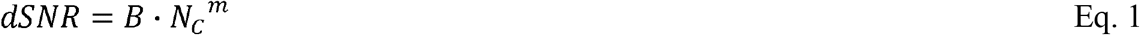

where *B* is an arbitrary constant, *m* is the slope (varying 0.34-0.43), and *N*_*C*_ is the channel count. The empirically determined growth (*m* = 0.34 to 0.43) could be less than the ideal of *m* = 0.5 because of behavioral confounders or violations of noise assumptions (independent, identically distributed, i.i.d., gaussian noise; see Methods).

### Translation of a finger iBCI to virtual quadcopter control

While an obvious clinical application of a finger iBCI is to restore fine motor control for a robotic arm^8^ or to reanimate the native limb^14^, a finger iBCI system could also be an intuitive approach to controlling multiple simultaneous digital endpoints, extending the functionality of 2D cursor control^10^. Another application for multiple-DOF finger control is video gaming, aimed at enabling people with disabilities to participate in this activity with others. To this end, each finger movement was mapped to a DOF for control of a virtual quadcopter (Fig. 4a). Unlike a previous implementation of a flight simulator^29^, the finger positions were mapped directly to velocity control of the quadcopter and not transformed into “quadcopter space” during retraining. Mapping finger positions to velocity control could also allow a general-purpose control paradigm for a variety of games. The only task-specific adaptation was to apply a low-level velocity back to neutral when the fingers were within 10% (of the total range of motion) of the neutral position. This kept the fingers in the neutral position unless the participant deliberately moved them. The positions of the fingers were visible in the bottom left portion of the screen with annotations indicating the neutral position of each finger and the cardinal directions for the thumb movements (Fig. 4b, top panel).

**Figure 4.**
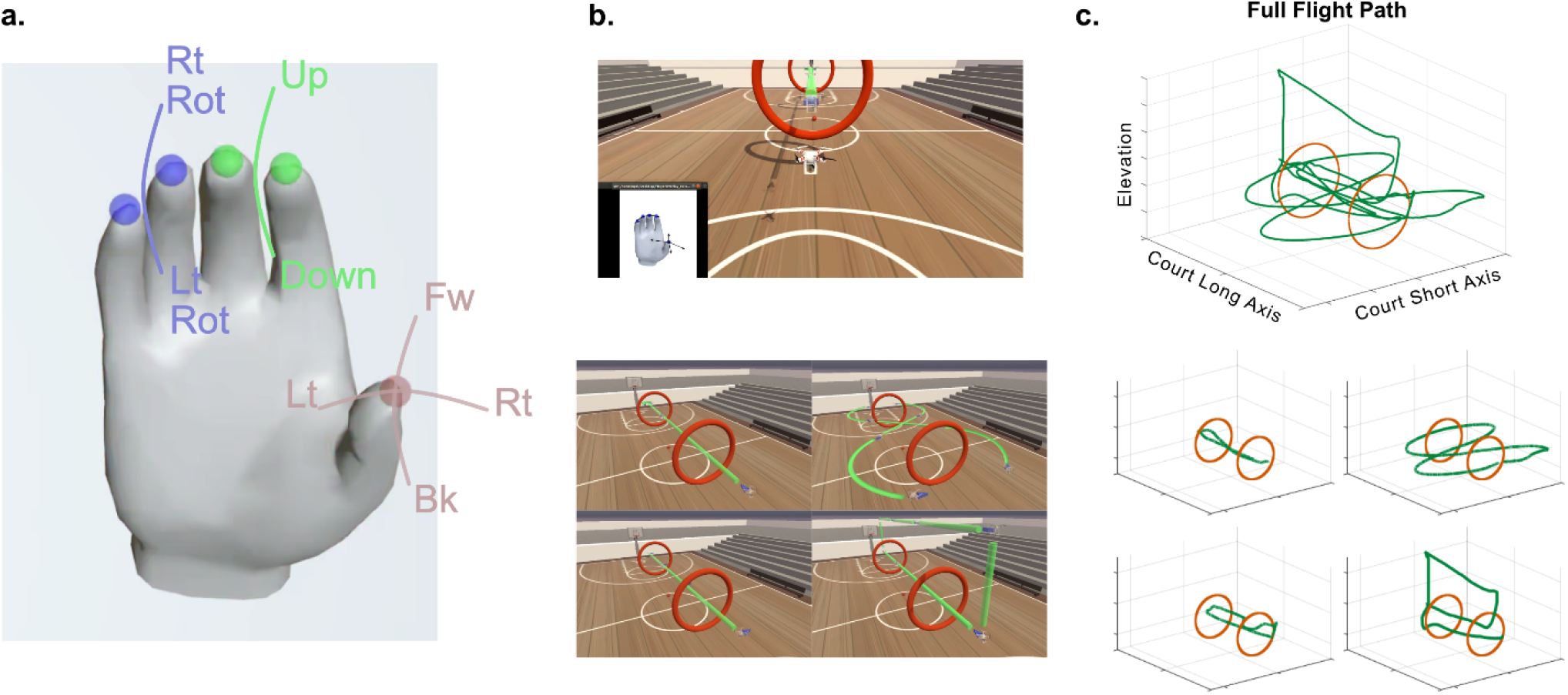
Finger intracortical brain-computer interface translated to virtual quadcopter control. **a**, Mapping finger position to quadcopter velocities. The thumb position is mapped to forward (Fw), backward (Bk), left (Lt), and right (Rt) translation velocity. The index-middle finger group position is mapped to velocities directed up and down in elevation. The position of the ring-small finger group is mapped to right rotation (Rt Rot) and left rotation (Lt Rot) velocities. **b (top pane)**, The layout for quadcopter control showing the virtual quadcopter in the center of the screen. A visualization of the hand indicating the neutral points for the finger groups and cardinal directions of the thumb is also visible. The rings are seen in the center of the display, and the green straight line indicates the trajectory the quadcopter is to follow along the obstacle course. **b (bottom pane)**, The quadcopter obstacle course demonstrates the 4-DOF control required to complete the 4.5-lap obstacle course. The top-left path requires the quadcopter to move forward, turn around, and move forward through the same rings to return to the starting point (1 lap). The top-right path requires the participant to simultaneously move forward and turn to complete 2 “figure-8” paths around the rings and back to the starting point (1 lap). The bottom-left path requires him to move left through both rings, stop, and then move right back through the rings (1 lap). The bottom-right path requires moving forward through the rings, increasing the elevation, moving backward over top of the rings, decreasing elevation, and then moving forward through both rings to the ending point (1.5 laps). **c (top pane)**, An exemplary full-flight path during a block of the obstacle course. **c (bottom pane)**, The flight path is separated into laps corresponding to the planned flight path for each lap in **b (bottom pane)**.

To demonstrate all the possible 4-DOF movements, an obstacle course was created (Fig. 4b, bottom panel) where each course segment could demonstrate at least one of the movements. On a single day of testing, the participant controlled the quadcopter over the complete obstacle course on 12 blocks with an average block time of 222 s and a standard deviation of 45 s. An exemplary block, completed in 163 s, is shown in Movie 5 with the flight path depicted in Fig. 4c. Since all fingers could be simultaneously decoded, multiple quadcopter movements could be combined with multiple finger movements, such as when the quadcopter moves forward and turns during the figure-8 segment of the obstacle course. Furthermore, since the finger positions lie along a continuum, a range of velocities can be provided for quadcopter control, which allows for high-velocity movements to cover large distances or low-velocity movements for fine adjustments.

While the obstacle course demonstrates 4-DOF control, the quadcopter was also tested in a less scripted, free-form task in which the participant was instructed to fly the quadcopter through randomly appearing rings (timeout every 20 s). This task illustrates reaction time, corrective maneuverability, and the ability to combine simultaneous DOF. After training the decoder, the participant was asked to fly through the rings. Over 10 min, he flew through 28 rings (2.8 rings/min); an illustrative segment from this session is given in Movie 6. Importantly, performance was impacted not only by decoding accuracy but also largely by behavioral factors, as even able-bodied operators using a unimanual quadcopter control might find the task challenging.

## Discussion

Motor BCIs have the potential to restore function for people with severe motor deficits from a variety of neurological diseases and injuries. While considerable work has advanced reaching/grasping for robotic arms and pointing/clicking for digital cursors^10,14-17^, continuous control of finger movements in human participants – necessary for fine motor movements – has received much less attention. In this work, real-time, closed-loop, 4D dexterous decoding demonstrated control of 3 highly-individuated finger groups with 2D thumb movements and allowed positioning of fingers at randomly selected targets within 1.58 ± 0.06 s (76 ± 2 targets/min) with peak performance reaching 1.30 ± 0.08 s (92 targets/min). This finger decoding BCI was mapped to control 3 digital effectors (with 1 effector moving in 2 dimensions) for high-performance, 4-DOF control of a virtual quadcopter. Although decoding an additional 2 DOF led to a nonlinear increase in the dimensionality of neural activity, training a decoder for an expanded set of movements did not degrade performance when testing the decoder on a smaller subset of movements where the neural representation was largely preserved. Finally, decoding accuracy, as measured by dSNR, was found to increase with input channel count at a sublinear rate for this channel count regime.

Simultaneously decoding 3 individuated finger groups with 2D thumb movements doubled the number of decoded DOF and achieved a similar level of performance as previous NHP finger-decoding work^22,23^. Acquisition times for the 2D task averaged around 1.3 s, and 98% trials completed and reached acquisition times as low as 0.84 s. This compares favorably to the 1.27 s acquisition time in NHPs, although the NHP task was a more complex random finger task.^23^

While dexterous decoding of 3 individualized finger groups with 4 DOF is a step toward fine motor control, decoding more DOF may be needed. When transitioning from 2D to 4D decoding, a greater than 2-fold increased dimensionality of neural activity was observed. The nonlinear increase in dimensionality supports the hypothesis that neural encoding for combinations of finger movements is not an exact linear superposition of the individual components, and this nonlinearity has been hypothesized by others^22,30^. In a recent study examining how simultaneous finger movements are encoded^30^, the dimensionally-reduced neural activity (in vector space) was found to be similar in angle but reduced in amplitude when compared to the sum of the individual component movements.

It is becoming increasingly evident that multiple effectors and DOF may be represented within the same neural population in motor cortex^18,27^. We demonstrated that decoder performance did not substantially decline when using a higher-DOF decoder to decode a lower-DOF closed-loop task, and the neural representation of finger movements appeared similar regardless of how many DOF could be actively controlled. How motor cortex represents movements of multiple effectors and limbs is an area of active investigation, with high-DOF representations emerging as a general principle^27^.

Although motor cortex can represent both 4D and 2D movements, acquisition times on the 4D task were longer than acquisition times on the 2D task. Many factors could lead to a slower performance on the 4D task (such as the increased difficulty inherent in the task), and our participant reported that keeping fingers stationary on the targets was challenging. Decoding nonzero velocities when trying to hold a finger stationary (i.e., signal-independent noise) caused the decoded finger position to drift off target. Several nonlinear approaches have been developed that could help mitigate this issue^23,31,32^, such as using hidden Markov models to estimate movement and stationary periods^32^. These approaches may be increasingly important with higher numbers of independently decoded effectors.

We also introduced a surrogate for decoding accuracy, dSNR, and found that dSNR was not saturating at our current channel count of 192 channels. This suggests that BCI systems are likely to improve with higher channel count systems. Improved decoding accuracy could not only lead to improvement in current BCI applications (i.e., cursor control, robotic arms, and finger BCI systems) but also allow for expanded functionality and new, more sophisticated applications.

The finger iBCI system developed in this work was used to control a high-performance virtual quadcopter. Compared with a state-of-the-art electroencephalographic (EEG)-controlled quadcopter that navigated through 3.1 rings in 4 minutes^33^, our system allowed navigation through or around 18 rings – at peak performance – in less than 3 minutes on a similar flight path, a more than sixfold increase in performance. The system was also capable of spontaneous free-form flight through randomly appearing rings. This level of performance demonstrates the feasibility using iBCI systems to control video games, virtual reality, or other digital interfaces. Importantly, these systems may address the unmet needs of people with paralysis for peer support, leisure activities, and sporting activities^34^, and may also allow more functionality compared to other assistive technologies such as mouth styluses or eye tracking software.

Although high performance was achieved using this decoding system, potential improvements to increase the likelihood of clinical adoption include reducing the calibration times and increasing the robustness to neural instabilities. Several approaches could be applied to this decoding system, including rapid decoder calibration^35^, training decoders using a long history of previously recorded data^36^, adaptive decoders using task knowledge^37,38^, and algorithms that perform dimensionality reduction to a stable manifold followed by realignment^39,40^.

In summary, we developed a high-performance finger iBCI system that could be mapped to multiple digital endpoints to allow people with paralysis to interact with others socially through video game play, virtual reality, or other digital connections.

## Acknowledgements

The authors thank the participant T5 for his generously volunteered time and effort as part of the BrainGate2 clinical trial. We acknowledge Krishna Shenoy for his inspiration and effort toward creating an environment that spawned this work. We thank David Sussillo for his thoughtful discussions, Michael K. Lim for support from the Department of Neurosurgery at Stanford University, and Beverly Davis, Kathy Tsou, and Sandrin Kosasih for administrative support.

This work was supported by the Office of Research and Development, Rehabilitation R&D Service, Department of Veterans Affairs (N2864C, A2295R); Wu Tsai Neurosciences Institute; Howard Hughes Medical Institute; Larry and Pamela Garlick; Samuel and Betsy Reeves; Sons Foundation Collaboration on the Global Brain 543045; NIH-NIDCD R01-DC014034, and NIH-NIDCD U01-DC017844.

*The contents do not represent the views of the Department of Veterans Affairs or the U.S. Government.

CAUTION: Investigational Device. Limited by Federal Law to Investigational Use

## Author Contributions

Conceptualization, M.S.W., N.P.S., D.T.A., J.M.H.; Methodology, M.S.W., N.P.S., D.T.A., F.R.W., J.M.H.; Software, M.S.W, N.P.S., D.T.A., F.R.W.; Validation, Software, M.S.W., N.P.S., D.T.A., F.R.W.; Formal analysis, M.S.W.; Investigation, M.S.W., N.P.S., D.T.A., N.V.H., R.M.J., F.B.K., F.R.W., J.M.H.; Writing – original draft, M.S.W.; Writing – review & editing, M.S.W., N.P.S., D.T.A., N.V.H., R.M.J., F.B.K., L.R.H., F.R.W., J.M.H.; Supervision, M.S.W., F.R.W., J.M.H.; Project administration, L.R.H., J.M.H.; Funding acquisition, L.R.H., J.M.H.

## Methods

### Clinical trial and participant

The participant, T5, was enrolled as a participant in the BrainGate2 Neural Interface System clinical trial (NCT00912041, registered June 3, 2009) with an IDE from the FDA (IDE #G090003). This study was approved by the Institutional Review Board (IRB) of Stanford University (protocol #20804) and the Mass General Brigham IRB (protocol #2009P000505). All research was performed while following all relevant regulations.

The participant, T5, was a 69-year-old right-handed man with C4 AIS C spinal cord injury. In 2016, 2 96-channel microelectrode arrays (Neuroport arrays with 1.5-mm electrode length; Blackrock Microsystems, Salt Lake City, UT) were placed in the anatomically identified hand ‘knob’ area of the left precentral gyrus. Detailed array locations are depicted on an MRI-reconstructed graphic in the Extended Data Fig. 1a (from Deo et al.^31^ in Extended Data Fig. 1a). Below the level of injury, T5 had very low amplitude movements that consisted primarily of muscle twitching. Tuning of the microelectrode arrays to these finger movements was confirmed (Extended Data Fig. 1b-c).

### Participant sessions

A total of 9 sessions of 2-5 h/session between March and August of 2023 were used to demonstrate online, closed-loop finger decoding and quadcopter control. The participant laid flat in bed with the monitor positioned above and slightly to his left so that he could keep his neck in the neutral position. Data were collected in roughly 1–10-min blocks. In between blocks, T5 was encouraged to rest as desired. Descriptions of the data collection sessions are shown in Extended Data Table 3.

### Finger tasks

A virtual finger display was developed in Unity (2021.3.9f1) that allows control of virtual fingers. The thumb was programmed to allow movement in 2 dimensions (flexion/extension and abduction/adduction), the index-middle fingers were grouped to move together within a 1-dimensional flexion/extension arc, and the ring-small fingers were grouped together to move in a 1-dimensional flexion/extension arc. By supplying a value between 0 and 1 for each of the 4 DOF, the finger position could be placed at continuously varying positions between full flexion and extension or abduction and adduction. Finger position values were set to follow pre-programmed trajectories during the open-loop blocks and were specified by the decoding algorithm during the closed-loop blocks.

#### Open-loop finger task

Center-out-and-back trials were paired together. On the “center-out” trials, one of the 3 finger groups was randomly chosen (or 2 finger groups when training the 2D decoder) to move from the neutral position to either full flexion or full extension in 2 s and then hold for 1 s. The participant was asked to attempt movement of his fingers in sync with the virtual fingers following a smoothly varying trajectory. On the “back” trial, the previously flexed or extended finger group would move back toward the neutral position and then hold for 1 s. Rest trials without finger movement were also included. To allow comparison with previous and future finger work^19,20,30^, finger movements were also classified using neural activity over long time windows typically used in classification (2 s) and short time windows typically used for closed-loop decoding (150 ms; Extended Data Fig. 1).

#### Closed-loop 2D finger tasks

The closed-loop 2-finger task was used for both training and testing the decoding algorithm. In this task, the participant controlled 2 simultaneous finger groups within a 1-dimensional arc: the thumb and index-middle group. On paired trials, the participant was cued to simultaneously move the finger groups from a center “neutral” position toward random targets within the active range of motion. Once reaching the target, all fingers were required to be within the target for 500 ms for the trial to be successfully completed. On the subsequent trial, targets were placed back at the center. The target width was 20% of the range of motion, and the trial timeout time was 10 s.

#### Closed-loop 4D finger tasks

There were several 4D finger tasks used for training and testing the decoding algorithm. The most frequently tested 4D task, denoted 4T, allowed the participant to simultaneously control 3 finger groups: thumb with 2D movements of flexion/extension and abduction/adduction, the index-middle group with 1D movements of flexion/extension, and the ring-small group with 1D movements in flexion/extension. In the first of paired trials, 2 new random targets would appear for 2 randomly selected finger groups, and the participant would be cued to move the fingers to the targets while keeping the third finger group stationary within its original central position target. The trial was completed successfully, if all 3 finger groups were within their respective targets for 500 ms before a 10 s trial timeout. On the second of 2 paired trials, all targets would return to the center position, prompting the 2 moving fingers from the previous trial to return to center targets. A similar task (Extended Data Fig. 5a), had only 1 new target/trial. Finally, when training the quadcopter, a closed-loop random finger task was used, where 2 new random targets per trial appeared in the active range of motion for the finger groups, i.e., there were no paired center-out-back trials, and each trial was independent of the previous.

The most used task for training, denoted T_TRAIN_, was a 4-DOF task similar to 4T above with several key differences so that intended decoder movements could be accurately inferred from a poorly/partially trained decoder. First, at the end of each trial, the positions of the fingers would return to the center position, which prevented fingers from becoming permanently stuck in flexion or extension. When 2 new targets were presented on a trial, the finger without a new target was artificially held fixed in the center position so that the participant could focus on only 2 finger groups per trial. The required hold time to successfully complete a trial was lengthened to 1.5 s to provide more training data when trying to steady the fingers, and trial timeout was reduced to 5 s so that the participant would not decrease his effort at the end of a longer trial. Finally, every other trial held the targets in the center position and the virtual fingers were fixed in place to provide an abundant amount of data where the participant was trying to remain stationary on the targets.

### Quadcopter tasks

To demonstrate the utility of closed-loop, online dexterous finger decoding in an applied task, finger control was mapped to 4D control of a virtual quadcopter. Specifically, the finger positions were mapped to a velocity-control paradigm, as shown in Fig. 4a. A physics-based quadcopter environment used the Microsoft AirSim plugin^41^ as a quadcopter simulator in Unity (2019.3.12f1). Two main tasks were developed to test this control: the quadcopter obstacle course that demonstrates control with all 4 DOF and the random ring acquisition task in which the participant demonstrates spontaneous control using multiple DOF at the same time. The participant was given time to become comfortable with the control paradigm in some preliminary sessions and was then evaluated on the obstacle course and random ring task for 1 day each.

#### Quadcopter obstacle course

A virtual basketball court was created in Unity with two large rings placed along the long-axis of the basketball court (see Fig. 4b). To demonstrate control of all 4 DOF, a path through and around the rings was designed (Fig. 4b). The participant was instructed to complete all segments of the obstacle course as quickly and accurately as possible, but no penalty was assessed for not staying exactly on course. During the day, he completed 12 blocks, and the decoder was retrained periodically to optimize performance.

#### Random ring acquisition

On 1 day of testing, only 1 ring was displayed, which was randomly generated both in its location in space and orientation, and the participant navigated the quadcopter through these random rings. The rate of ring acquisition during the first 10 min was calculated, and a movie of a representative time segment is included.

### BCI rig and front-end signal processing

The BCI rig was set up in 3 distinct configurations as our lab transitioned from an older analog setup to the newer digital setup. In the first setup used until 7/10/2023, 2 patient cables were connected to the transcutaneous pedestals, which were routed to the Neural Signal Front End Amplifier (Blackrock Neurotech, Salt Lake City, Utah) where the raw voltage was bandpass filtered (0.3 Hz first-order high-pass and 7.5 kHz third-order low-pass), sampled at 30 kHz with 250 nV resolution, converted to an optical signal, and then sent to the Neural Signal Processor^42^. From 7/26/2023 and later, 2 Neuroplex E headstages were connected to 2 transcutaneous pedestals, and the signal was analog filtered and sampled at the headstage and then sent to the digital hub via a micro-HDMI cable. At the digital hub, the signal was converted to an optical signal transmitted via optical cable to the Neural Signal Processor. In the final mixed configuration, used between 7/12 and 7/24/2023, the setup using the analog patient cable was connected to the anterior pedestal, and a Neuroplex E headstage with its subsequent configuration was connected to the posterior pedestal.

The Neural Signal Processor sends the digital signal to a SuperLogics machine running Simulink Real-Time (v2019, Mathworks, Natick, MA). For most sessions, common average referencing (CAR) was used to reduce the electrical noise on the input channels. However, during both quadcopter sessions (sessions 8 and 9), CAR was switched to linear regression referencing to predict a reference from the combined input channels that is subtracted from each input channel^43^. The signals then passed through a 250-Hz digital high-pass filter. The data were binned into 50-ms windows. The sum of the squared magnitude was calculated for each window. This signal was denoted as spike-band power (SBP). Every 50 ms, UDP packets of neural features are communicated to a Linux computer running Ubuntu with Python (v3.7.11), PyTorch (v1.12.1, https://pytorch.org/), and Redis (v7.02), where the neural features pass into the decoding algorithm. The entire system was interfaced with an additional Windows computer running Matlab (v2019, Mathworks, Natick, MA) that was interfaced with the system to stop and start experimental runs during sessions.

### Decoding algorithm

The decoding algorithm presented by Willsey et al.^23^ was adapted for this work. The algorithm is a shallow-layer feed-forward neural network with an initial time-feature learning layer implemented as a scalar product of historical time bins and learned weights. A rectified linear unit (ReLU) was used as the non-linearity after the convolutional layer and each linear layer except for the last linear layer. The input ***Y***_*IN*_ was an ***E***_*N*_ x 3 input matrix, where ***E***_*N*_ is the number of electrodes (192) and 3 represents the 3 most recent 50-ms bins. The time feature learning layer converts 3 50-ms bins into 16 learned features using weights that are shared across all input channels. The output was flattened and then passed through 4 fully connected layers.

The intermediate outputs were highly regularized with batch normalization (batchnorm)^44^ and 50% drop out. The output variable, 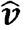, represents an array of decoded finger velocities, that if ideally trained, would be normalized with zero mean with unit variance. However, an empirical mean value and standard deviation were subsequently calculated from the training data set, which were used to normalize 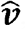, and then an empirically-tuned gain was applied to the decoded finger velocities.

In a change from Willsey et al.^23^, to reduce the ability of the neural network to produce velocities with non-zero means, the final linear layer was changed to disallow an affine output and the final batchnorm layer was not allowed to learn a bias. Furthermore, during training and testing, the final batchnorm was not allowed to apply a mean correction, as only a variance correction was allowed. The purpose of these changes was to penalize the preceding algorithmic blocks during training if the decoded signal had a non-zero mean.

### Closed-loop decoding software

The SBP was imported to a script that calculated 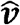 from the input data (3 time bins, 192 channels). The signal 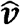 was normalized using the values calculated during training and the empirically tuned gain was also applied. No smoothing was applied. The positions of the fingers were updated at each time step using the velocities.

When the positions of the virtual fingers were used to control the quadcopter, “gravity” was applied to the fingers when the fingers were near the neutral position so that the fingers did not appear to jitter when the intention was to hold them steady. Specifically, when the fingers were within 10% of the range of motion of the neutral position, a position-independent, constant, low-amplitude value was added to the decoded velocity of the finger to bias the velocity toward the neutral position. Decoded velocities were scaled to a maximum of ±10 m/s and ±90 deg/s for linear and rotational velocities, and each DOF was tuned empirically with gain values equal to 0.6 for thumb flexion/extension, 0.8 for thumb abduction/adduction, 0.4 for index-middle flexion/extension, and 0.6 for ring-small flexion/extension.

### Offline algorithm training

The algorithm was trained on a combination of open- and closed-loop trials. For each day, the algorithm was trained on 2 blocks of 100 open-loop trials. The SBP data were organized into batches of 64×256×3 (64 randomly selected time steps, 256 input channels, 3 previous time bins adjacent in time to the current time step). The velocities of the virtual fingers during the open-loop block were normalized by the standard deviation of the velocity and then multiplied by 0.2 (referred to as `stretchFactor`), as a reduced amplitude of open-loop finger velocities was previously observed to yield a better offline fit^23^. Thus, the finger velocities used for training were (in pseudocode):

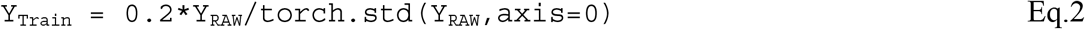

where Y_RAW_ was an array of *d* finger velocities for *N* training samples (*N*×*d*).

The algorithm (Extended Data Fig. 2) was initialized using the Kaiming initialization method^45^. The neural network minimized the mean-squared error (torch.nn.MSELoss) between the actual finger velocities during open-loop training and the algorithm output using Adam optimization algorithm^46^ (torch.optim.Adam). The optimizer was used with a learning rate of 10^−4^ and weight decay of 10^−2^ (parameters: lr=1e-4, weight_decay=1e-2), and the algorithm was trained over 10 epochs.

After training the algorithm, a mean offset value, *μ*_*offset*_, was calculated (Eqs. 3 and 4) to give the output of the neural network algorithm the same mean as the unnormalized training data.

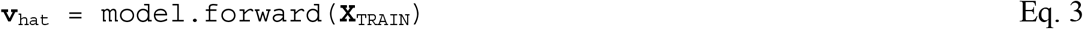

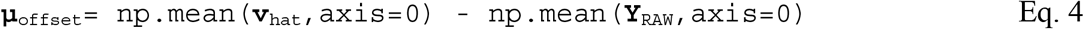

The variable **v**_hat_ is the (*N*×*d*) output tensor after applying the trained neural network algorithm (`model)`, **X**_TRAIN_ is a *N*×*E*_*N*_ *×*3 tensor array for the *N* training time steps, *E*_*N*_ input channels, and 3 preceding time steps to the current time step. The variable **v**_hat_ was converted to a numpy array and input to Eq. 3, and Y_RAW_ was defined in Eq. 2. A gain for the algorithm output, *G*, was calculated to normalize the standard deviation of the algorithm output and further reduce the output amplitude by a factor of 3 (which was empirically determined):

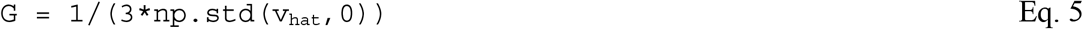

Thus, when running the algorithm for online, closed-loop decoding, the output for each time step was adjusted according to Eq. 6:

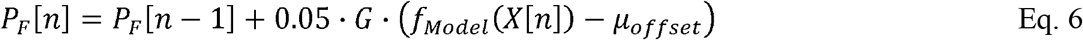

where *P*_*F*_*[n]* denotes the position of *d* finger groups at time step *n*.

### ReFIT training

After the offline algorithm training, the online, closed-loop sessions were performed. After a closed-loop session, the adapted recalibrated feedback intention-trained (ReFIT) algorithm^23,47^ was used to update the parameters of the neural network. Similar to above, SBP data were organized into 64×256×3 batches, with the 64 time steps randomly selected. The corresponding finger velocities used for training were assigned a value equal to the decoded velocity when the velocity is pointed toward the target, and the sign is inverted when the velocity is directed away from the target (Eq. 7):

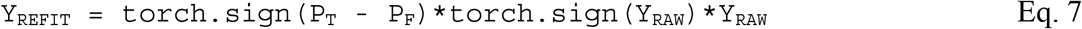

where P_T_ is the position of the target, P_F_ is the position of the fingers, and Y_RAW_ was an array of *d* finger velocities for *N* training samples (*N*×*d*). Similar to offline training, the velocities were then scaled by the standard deviation and a `stretchFactor` of 1.3 (promoting higher velocities toward the target), except when the finger positions lie within the target when `stretchFactor` divides the value of *Y*_*REFIT*_ (promoting lower velocities). These steps were implemented by executing the 2 consecutive lines of pseudocode given in Eqs. 8-9:

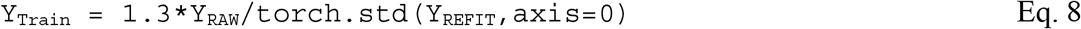

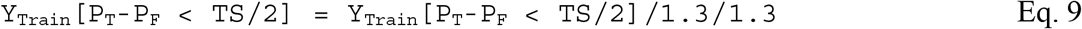

Of note, dividing twice by `1.3` in Eq. 9 is required to undo the `1.3` multiplication factor in Eq. 8. Starting with the same parameters for the neural network algorithm used during the online session, the Adam optimization algorithm (lr=1e-4, weight_decay=1e-2) was applied and trained over 500 additional iterations. A new value for *μ*_*offset*_ was calculated according to Eq. 8 using `np.median` instead of `np.mean`, and *G* was calculated as before in Eq. 5.

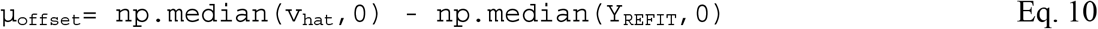

When running the algorithm online, the finger positions were again updated according to Eq. 6.

### Training protocols for the 4D decoder

After the algorithm parameters were trained from the open-loop session, closed-loop control using T_TRAIN_, which was easier to control with a suboptimal decoder, was used until approximately 80% of trials were completed. Then the 3-finger task, 4T, was used for 50 additional trials. After each closed-loop session, the algorithm parameters were updated according to the above section (**ReFIT training**).

As a control to understand how neural instabilities^48^ could affect decoding performance, the stability of the 4D decoder was evaluated during 2 research sessions by training an initial decoder, fixing the parameters, and using this fixed decoder on consecutive blocks until trials could not be reliably completed. This occurred after 20 minutes (5 blocks) on the first day and 53 minutes (11 blocks) on the second. On the first day, the decoder was re-trained to demonstrate recovery of performance with re-training (Extended Data Fig. 5b).

On occasion the decoder was trained but the parameters required updating either to improve performance from an instability or for a fair comparison with another decoder. When this was required, a combination of T_TRAIN_ and 4T were used. The training of each decoder used in closed-loop sessions is described in Extended Data Table 4.

### Training protocols for the 2D decoder

The 2D finger decoder was trained with open-loop sessions first and then with closed-loop sessions, like the 4D decoder. Unlike the 4D decoder, the 2-finger task for the 2D decoder was the only task performed. Furthermore, on some occasions, the 2D decoder was trained until 100% of trials were completed successfully, and on other occasions training was continued even after 100% of trials were completed. The training of these decoders is also described in Extended Data Table 4.

### Online performance metrics

Various metrics were calculated to characterize online performance. Trials in which fingers started within a “new” target were excluded from the analysis. Performance metrics were only calculated from successful trials. The acquisition time was defined as the time from the start of the trial to when the fingers had successfully held on each of the targets for the required hold time of 500 ms subtracted by 500 ms. The time to target was defined as the time from the start of the trial to when all fingers reached the target (and did not require all fingers to be on the target simultaneously). The orbiting time was the acquisition time subtracted by the time to target. Targets per minute was the number of new targets per trial divided by the mean acquisition time (excluding hold time). Successfully completing the tasks required completing the task within 10 s (i.e., acquisition time of 9.5 s). For completeness, the path length efficiency was calculated, although a metric perhaps more suitable for single-effector *d*-dimension control. Path length for each trial was calculated according to Eq. 11:

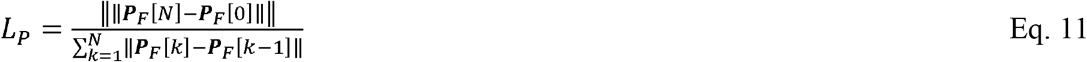

Where ∥·∥denotes the L_2_ norm, *L*_*p*_ is the path length efficiency for that trial, *N*+1 is the elapsed samples until all fingers are on the target, and ***P***_*F*_ is a vector of the positions of all *d* fingers. Thus, *L*_*p*_ close to unity implies the fingers traveled a direct path to the targets, and a value close to zero implies a circuitous path to the target. Finally, when calculating the throughput in bits per second (bps), *T*_*bps*_, the same adaption of Fitt’s law for fingers developed in Willsey et al.^23^ was used and is repeated in Eq. 12:

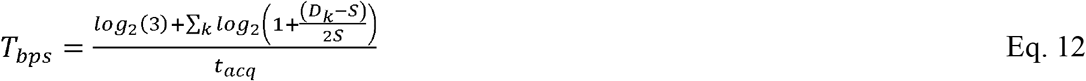

where *k* indexes through all 3 finger groups, Dk is the distance of the *k*-th to the target, *S* is the circular target radius, and *t*_*acq*_ is the target acquisition time. *T*_*bps*_ was then averaged over all trials. For this calculation, the only trials included are those for which all fingers begin a distance from the center of the target that is greater than twice the target radius. In an adaption to the approach in Willsey et al.,^23^ log_2_(3) bits was added on each trial to account for the information needed to convey which finger is stationary. An alternative approach would be to not add these additional bits and allow the *T*_*bps*_ to decrease with the understanding that the total corpus of targets is higher.

To illustrate how decoded finger movements could be discriminated, the mean decoded velocity was calculated during single-finger movements. This analysis is shown in Fig. 1f. Four blocks of the 4D task with 1 new target/tr (Extended Data Fig. 5a) were used for this analysis. On each trial, the mean velocity of all fingers was calculated during the ‘Go’ period (200-700 ms after trial start) and normalized by the mean value of the finger group with the highest mean value (which was the cued finger).

### Offline analyses

The offline analyses were conducted in Python (v3.9.12) using a Jupyter notebook (https://jupyter.org/) and in Matlab (v2022a, Mathworks, Natick, MA). The following python packages were used: scipy (v1.7.3), torch (v1.12.0), torchvision (v0.13.0), numpy (v1.21.5), matplotlib (v3.5.3), PIL (v9.0.1), sklearn (v1.0.2).

### Statistical analysis

All statistical comparisons used a two-sample, two-tailed *t*-test in Matlab using the function: *ttest2*.*m*.

#### Confusion matrices

The fingers were classified during open-loop trials to relate the tuning of these arrays to other reports focusing on classification^19,20,30^. Classification over a 2-s movement window was used to illustrate performance over typical windows used for classification and over a shorter 150-ms window similar to windows used for closed-loop decoding. The open-loop data from a typical day, session 6, were used for this analysis (200 trials and 192 input channels of SBP). Both a 10-fold cross-validation and a linear discriminant analysis classifier that assumes a shared diagonal covariance matrix across conditions were used. The analysis was performed in Matlab 2022a using the functions: fitcdiscr.m (with ‘DiscrimType’ as ‘diaglinear’), crossval.m, kfoldLoss.m, and kfoldPredict.m.

#### Dimensionality

While there are numerous approaches to calculate dimensionality^49^, the participation ratio was used, which is roughly equivalent to the dimensions needed to capture 80% of the variance^11,50^. The ‘Go’ period during the trial, 200-700 ms after a new target appeared, was averaged for each condition. A *d* × 1 condition vector, ***C***_*DOF*_, was defined according to whether each respective DOF at the beginning of the trial needed to flex/abduct (+1), extend/adduct (−1), or remain stationary (0) to reach the targets. Thus, for 4D closed-loop decoding during the 4D finger task, ***C***_*DOF*_ =[1,1,0,−1]^*T*^ if the thumb needed to flex and abduct to reach the target, the index-middle group needed to remain on the target, and the ring-small group needed to extend.

The SBP for each electrode during the ‘Go’ period was z-scored and smoothed using scipy.ndimage.gaussian_filter1d with a sigma of 3 50-ms time bins. The data were then organized into a matrix, ***D***_*4d*_, that was *E*_*N*_×(*cN*_*M*_), where *E*_*N*_ is the integer 192 for the number of input channels, *c* is the integer 20 for the number of conditions in the 4D task with 2 new targets/trial, and *N*_*M*_ is the integer 10 for the number of 50-ms bins in the ‘Go’ period. Similar data matrices were calculated for the 2D task, ***D***_*2d*_, and for the 4D task with 1 target/trial, ***D***_*4d1t*_. The eigenvalues, *u*_*i*_, were then calculated for the cross-validated covariance matrix, 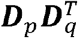 (where ***D***_*p*_ and ***D***_*q*_ were data matrices from 2 folds of the data). The participation ratio was calculated (Eq. 12).

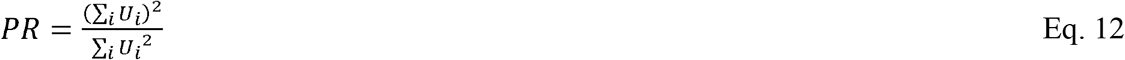

#### Analysis of the 2D and 4D decoders on the 2D task

To determine whether mapping changes when mapping neural activity to a 4D vs 2D task, decoders were trained on the 4D and 2D tasks as explained above and both of these decoders were used for the 2D task (thumb flexion/extension and index-middle flexion/extension). To compare these mappings, the 2D decoding algorithm was used to predict the velocities when using the 4D decoder in closed-loop trials, and the 4D algorithm was used to predict the closed-loop decoded velocities of the 2D algorithm. Fig. 2c illustrates velocities decoded online by the 2D decoder and predicted by the 4D decoder. To quantify the similarity between these signals, the normalized cross correlation, *r*_*n*_, function was calculated as defined below in Eq. 13,

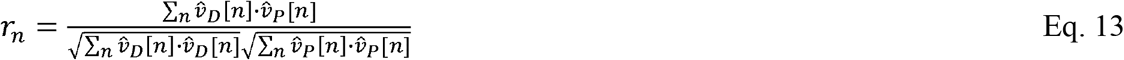

where 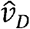 is the velocity decoded during the online block and 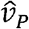 is the velocity predicted offline. The value for *r*_*n*_ was then averaged for both DOF. For paired blocks with the 4D and 2D decoders, the normalized cross correlation function was calculated, and these data are displayed in Fig. 2d.

#### Directional signal-to-noise ratio

While SNR metrics have been proposed for offline analyses, a vector-based SNR^51^ was adapted specifically for closed-loop decoding, denoted directional SNR (dSNR). In this formulation, ***v***[*n*] = [*v*_1_[*n*], *v*_2_[*n*], … *v*_*d*_[*n*]]^*T*^ is a normalized target vector, ∥***v***[*n*]∥ = 1, for *d* DOF with positive amplitudes for flexion/abduction and negative amplitudes for extension/abduction. Thus, in the 2D task, ***v****[n]*, at a given 50-ms time bin, *n*, is represented graphically in Fig. 3a, where, as an example, ***v*** = [0.707,0.707]^*T*^ is a 2-dimensional vector indicating that both fingers require flexion to reach the target. The array of *d* decoded/predicted finger velocities, 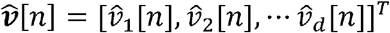, is assumed to be a time-varying, *d*-dimensional vector. This vector can be decomposed into orthogonal components, including a signal component, 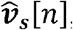, that is the projection of 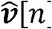along *v*[*n*], and a noise component, 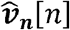, orthogonal to *v*[*n*], as graphically depicted in Fig. 3a for the 2D task. Using this formulation, dSNR is defined in Eq. 14:

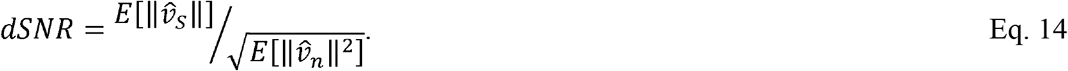

The value of dSNR was empirically calculated from closed-loop blocks of 2 and 3 decoded fingers (Extended Data Table 4) during the ‘Go’ period of the trials (200-700 ms after a new target was presented) before fingers were on their respective targets. To empirically calculate dSNR, the SBP data are divided into 6 folds: 5 training folds and 1 testing fold. To regularize the number of regressors (i.e., 192 channels) for linear regression, PCA decomposition was used (sklearn.decomposition.PCA) on the *n* 50-ms time bins by ***E***_*N*_ = 192 input channels (*n*×192) of SBP training data, ***X***_*TRAIN*_, to reduce the number of dimensions to an *n*×20 data set, 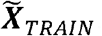. Using LinearRegression from sklearn.linear_model toolbox, a linear mapping is trained to map 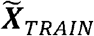 to the *N*×*d* training velocities, ***V***_*TRAIN*_., (i.e., ***v*** in Fig. 3a). These commands are represented with the pseudocode in Eqs. 15-18.

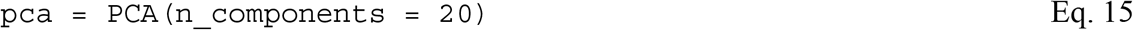

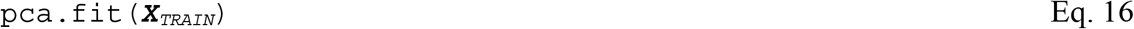

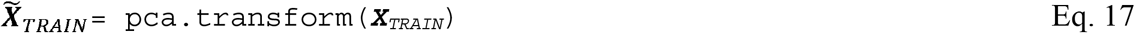

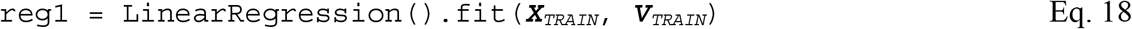

Finally, the predicted velocities, 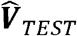, of the test data, 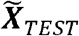 were determined from Eq. 19:

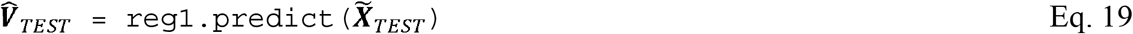

The predicted finger velocities, 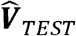, for the 2D decoder are shown in Fig. 3b. The magnitude of the signal component of the predicted velocity, 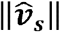 as in Fig 3a, was calculated from the dot product of 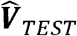 and ***V***_*TEST*_ according to Eq. 20:

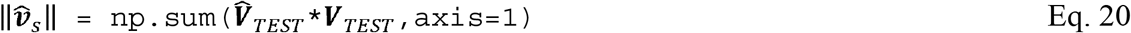

where 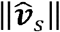 is length-*n* array for *n* time steps. To compute the noise component, 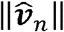, θ, the angle between 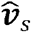 and ***v*** in Fig. 3a, and 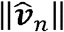 were calculated according to Eqs. 21 and 22. Finally, in Eq. 23, the value of `dSNR` was calculated.

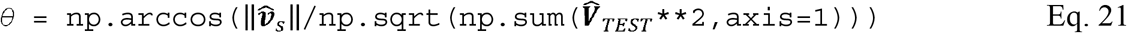

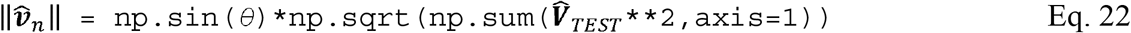

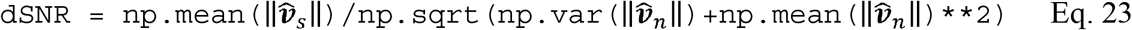

The value of `dSNR` was then averaged over all 6 folds. The data in 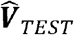 for all folds and all days are the population data, shown for the 2D decoder in Fig. 3b.

To calculate dSNR as a function of channel count, dSNR was calculated for an array of input channels, ***N***_*C*_*[k]*, indexed by *k* and ranging from 5 to the full *E*_*N*_ = 192 at a step size of *E*_*N*_/20. At each step, the value of dSNR was averaged over 25 iterations where at each iteration, ***N***_*C*_*[k]* random input channels were selected.

The empirical fit for the log of dSNR averaged over all days and log of ***N***_*C*_ was calculated using data from the highest 75% of values of ***N***_*C*_ and using numpy.linalg.lstsq for the empirical fit and sklearn.metrics.r2_score for the coefficient of determination, *R*^*2*^.

#### Theoretical SNR dependency on channel count

For a theoretical comparison for the dependency of dSNR on channel count, a 1D signal, *S*, measured independently on *N* channels was defined according to the form:

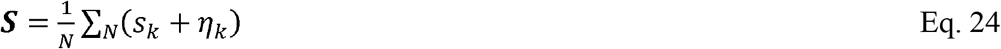

where *s*_*k*_ is the signal and *η*_*k*_ is i.i.d. Gaussian samples from the distribution *N*(0, *σ*). Assuming for simplicity and without loss of generality that *s*_*k*_ are equal, then Eq. 24 simplifies to:

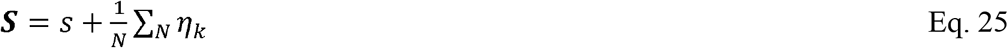

Thus, the value of the expected signal in Eq. 14 equals simply *s*. The expected square of the noise, *E*[***η***^2^], can be simplified to:

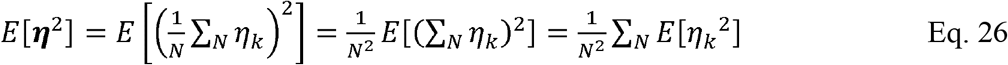

where the last equality follows since the terms η_k_ are independent. Finally,

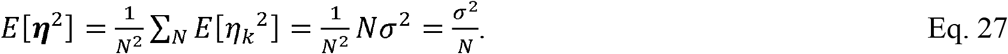

Thus, the SNR in this simplified case is:

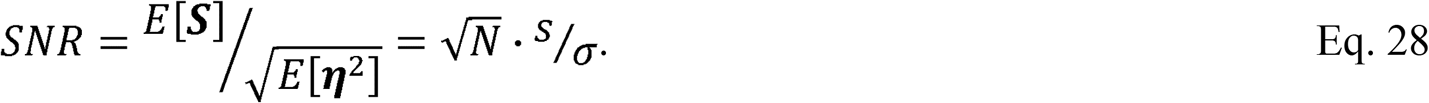

Thus, for our definition of SNR as defined in Eq. 14, the SNR increases proportionally with 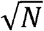 in this theoretical, simplified 1D formulation.

## Tables

**Extended Data Table 1:** Performance metrics for 4D finger decoding with 1 and 2 new targets per trial.

|  | 1 New target/trial | 2 New targets/trial |
| --- | --- | --- |
| Number of trials | 178 | 187 |
| Number of days | 1 | 1 |
| Acquisition time (ms) | 1380 ± 70 | 1660 ± 70 |
| Time to target (ms) | 890 ± 50 | 1260 ± 50 |
| Orbiting time (ms) | 480 ± 60 | 400 ± 50 |
| Targets per minute | 45 ± 6 | 74 ± 6 |
| Percent completed | 100% | 100% |
| Path length | 0.674 ± 0.014 | 0.607 ± 0.012 |

Statistical data are reported as mean ± standard error of the mean.

**Extended Data Table 2:** Performance metrics for finger decoding using 4D decoder on 2D task.

|  | 2D Decoder,<br>2D task | 4D Decoder,<br>4D task | 4D Decoder,<br>2D task |
| --- | --- | --- | --- |
| Number of trials | 233 | 284 | 329 |
| Number of days | 3 | 3 | 3 |
| Acquisition time (ms) | 1110 ± 50 | 1730 ± 70 | 1207 ± 40 |
| Time to target (ms) | 1040 ± 40 | 1210 ± 40 | 1060 ± 30 |
| Orbiting time (ms) | 60 ± 10 | 530 ± 50 | 150 ± 20 |
| Targets per minute | 111 ± 8 | 70 ± 3 | 101 ± 2 |
| Percent completed | 99.1% | 99.6% | 100% |
| Path length | 0.782 ± 0.010 | 0.558 ± 0.010 | 0.678 ± 0.009 |

Statistical data are reported as mean ± standard error of the mean.

**Extended Data Table 3:** Data sessions.

| Session no. | Date (post-implant day) | Description |
| --- | --- | --- |
| 1 | 3/9/2023 (2395) | Closed-loop finger decoding |
| 2 | 3/14/2023 (2400) | Closed-loop finger decoding |
| 3 | 3/16/2023 (2402) | Closed-loop finger decoding |
| 4 | 3/21/2023 (2407) | Closed-loop finger decoding |
| 5 | 3/23/2023 (2409) | Closed-loop finger decoding |
| 6 | 4/6/2023 (2423) | Closed-loop finger decoding |
| 7 | 4/13/2023 (2430) | Closed-loop finger decoding |
| 8 | 7/12/2023 (2520) | Quadcopter obstacle course |
| 9 | 8/28/2023 (2567) | Quadcopter random rings |

**Extended Data Table 4:**
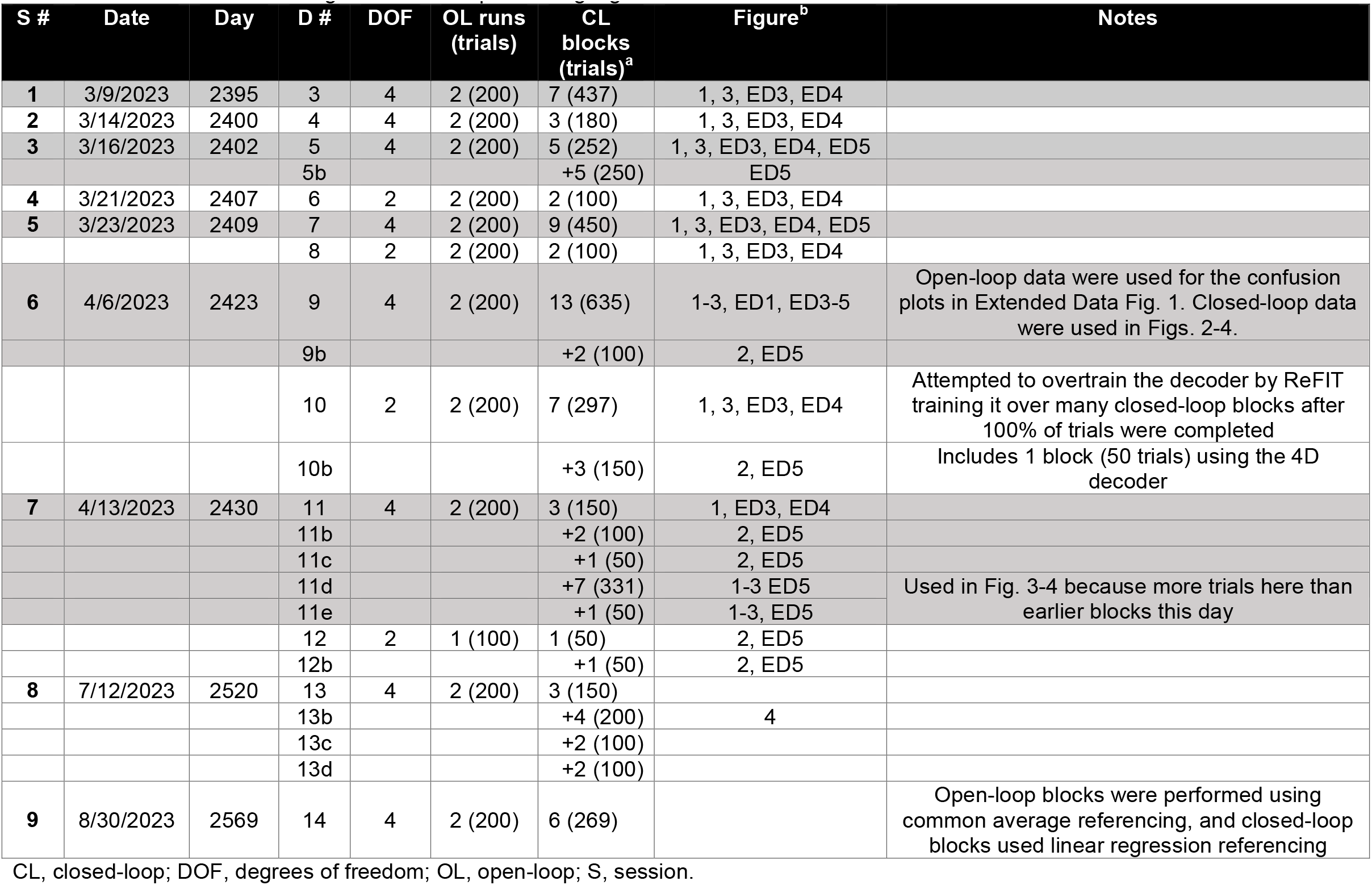
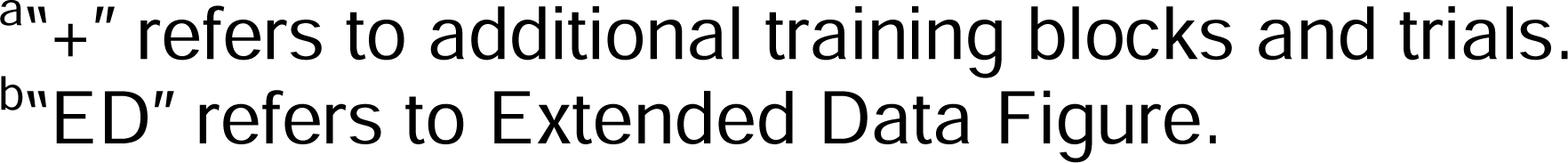
Training the closed-loop decoding algorithm.

| S # | Date | Day | D # | DOF | OL runs (trials) | CL blocks (trials)<sup>a</sup> | Figure<sup>b</sup> | Notes |
| --- | --- | --- | --- | --- | --- | --- | --- | --- |
| 1 | 3/9/2023 | 2395 | 3 | 4 | 2 (200) | 7 (437) | 1, 3, ED3, ED4 |  |
| 2 | 3/14/2023 | 2400 | 4 | 4 | 2 (200) | 3 (180) | 1, 3, ED3, ED4 |  |
| 3 | 3/16/2023 | 2402 | 5 | 4 | 2 (200) | 5 (252) | 1, 3, ED3, ED4, ED5 |  |
|  |  |  | 5b |  |  | +5 (250) | ED5 |  |
| 4 | 3/21/2023 | 2407 | 6 | 2 | 2 (200) | 2 (100) | 1, 3, ED3, ED4 |  |
| 5 | 3/23/2023 | 2409 | 7 | 4 | 2 (200) | 9 (450) | 1, 3, ED3, ED4, ED5 |  |
|  |  |  | 8 | 2 | 2 (200) | 2 (100) | 1, 3, ED3, ED4 |  |
| 6 | 4/6/2023 | 2423 | 9 | 4 | 2 (200) | 13 (635) | 1-3, ED1, ED3-5 | Open-loop data were used for the confusion plots in Extended Data Fig. 1. Closed-loop data were used in Figs. 2-4. |
|  |  |  | 9b |  |  | +2 (100) | 2, ED5 |  |
|  |  |  | 10 | 2 | 2 (200) | 7 (297) | 1, 3, ED3, ED4 | Attempted to overtrain the decoder by ReFIT training it over many closed-loop blocks after 100% of trials were completed |
|  |  |  | 10b |  |  | +3 (150) | 2, ED5 | Includes 1 block (50 trials) using the 4D decoder |
| 7 | 4/13/2023 | 2430 | 11 | 4 | 2 (200) | 3 (150) | 1, ED3, ED4 |  |
|  |  |  | 11b |  |  | +2 (100) | 2, ED5 |  |
|  |  |  | 11c |  |  | +1 (50) | 2, ED5 |  |
|  |  |  | 11d |  |  | +7 (331) | 1-3 ED5 | Used in Fig. 3-4 because more trials here than earlier blocks this day |
|  |  |  | 11e |  |  | +1 (50) | 1-3, ED5 |  |
|  |  |  | 12 | 2 | 1 (100) | 1 (50) | 2, ED5 |  |
|  |  |  | 12b |  |  | +1 (50) | 2, ED5 |  |
| 8 | 7/12/2023 | 2520 | 13 | 4 | 2 (200) | 3 (150) |  |  |
|  |  |  | 13b |  |  | +4 (200) | 4 |  |
|  |  |  | 13c |  |  | +2 (100) |  |  |
|  |  |  | 13d |  |  | +2 (100) |  |  |
| 9 | 8/30/2023 | 2569 | 14 | 4 | 2 (200) | 6 (269) |  | Open-loop blocks were performed using common average referencing, and closed-loop blocks used linear regression referencing |

CL, closed-loop; DOF, degrees of freedom; OL, open-loop; S, session.
<sup>a</sup>“+” refers to additional training blocks and trials. <sup>b</sup>“ED” refers to Extended Data Figure.

## Figures

**Extended Data Fig. 1.**
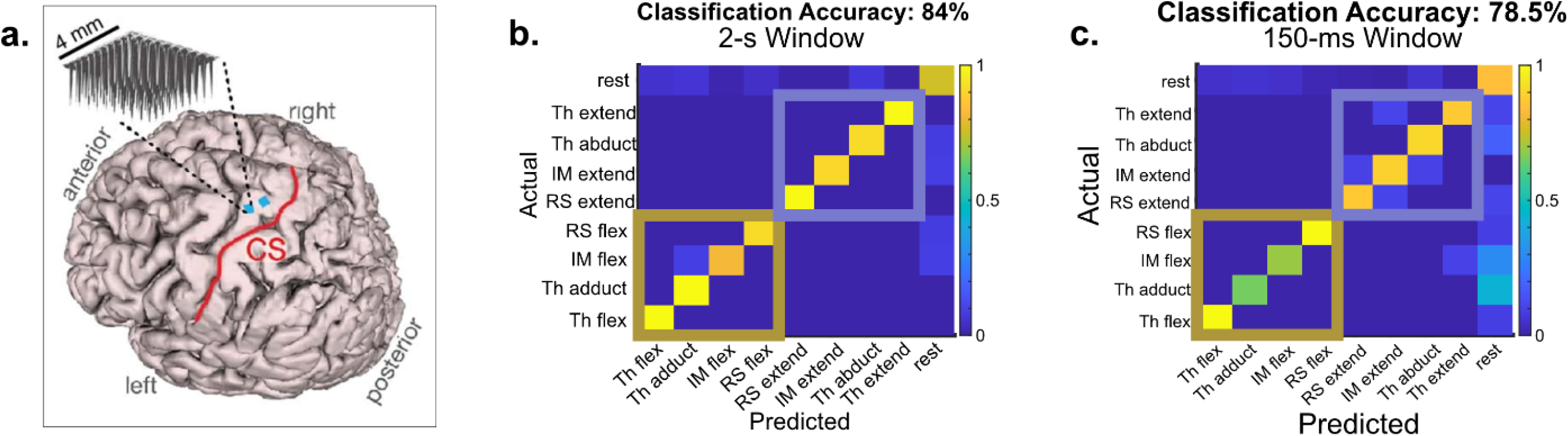
The intracortical brain-computer interface system for dexterous finger movements. **a**, MRI reconstruction of the participant’s brain with the implant locations depicted as blue squares. Two 96-channel silicon microelectrode arrays were placed in hand ‘knob’ area of left precentral gyrus in 2016. The red line indicates the central sulcus (CS). Panel from Deo et al. (2023)^31^. **b and c**, Confusion matrices showing the probability of correctly classifying attempted finger movements using 2 s, **b**, and 150-ms, **c**, windows from an offline analysis. IM, index-middle; RS, ring-small; TH, thumb.

**Extended Data Fig. 2.**
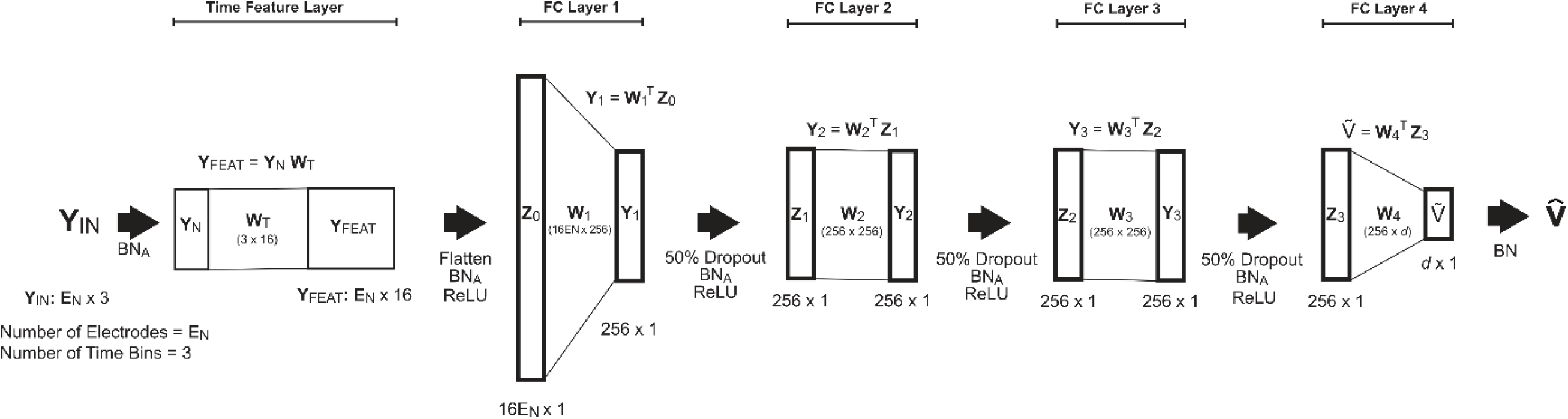
Decoding algorithm. The input ***Y***_*IN*_ is ***E***_*N*_ x 3 input matrix, where ***E***_*N*_ is the number of electrodes (192) and 3 represents the most recent 3 50-ms bins. The output variable, 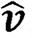, represents a normalized vector of each of *d* finger velocities. The actual decoded velocities were calculated by applying an empirically calculated mean value and gain value. Linear layers (***W***_*T*_, ***W***_*1*_-***W***_*3*_) included a learnable bias term except for the final linear layer, ***W***_*4*_, to reduce the magnitude of non-zero means. All instances of batchnorm, ***BN***_*A*_, were implemented with affine = *True* except for the final batchnorm, ***BN***, where affine = *False* in an attempt to reduce the reliance of the decoding algorithm on an offset correction from the final batchnorm block. During training mode batchnorm layer, ***BN***, did not correct for non-zero means or apply a mean correction to force the final linear layer, ***W***_*4*_, to learn an output signal with zero mean. BN, batchnorm; FC, fully connected; ReLU, rectified linear unit. Figure adapted from (Willsey et al., 2022)^23^.

**Extended Data Fig. 3.**
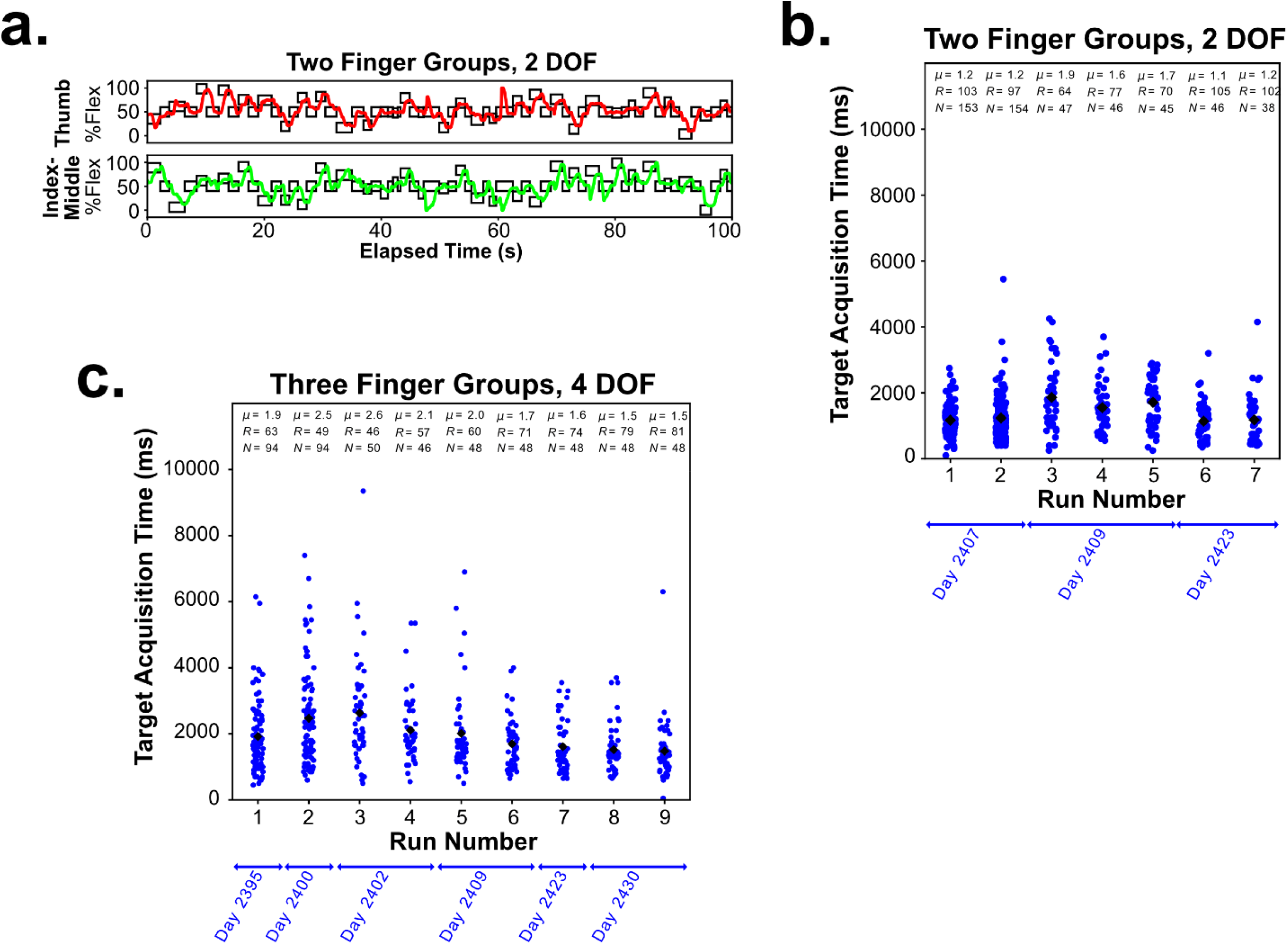
Closed-loop decoding for a 2- and 4-degree-of-freedom (DOF) finger task. **a**, A 100-s time segment of typical decoded movements is depicted for the 2-finger group, 2-DOF task. Trajectories are described as a percentage of flexion (%Flex). Distributions of target acquisition times (ms) for the 2-DOF task, **b**, and 4-DOF task, **c**, over multiple blocks of each task. Each dot corresponds to a trial, and the black diamond indicates the mean value. *N* denotes number of trials per block, R denotes the rate of targets acquired in targets/min, and *μ* denotes the mean value.

**Extended Data Fig. 4.**
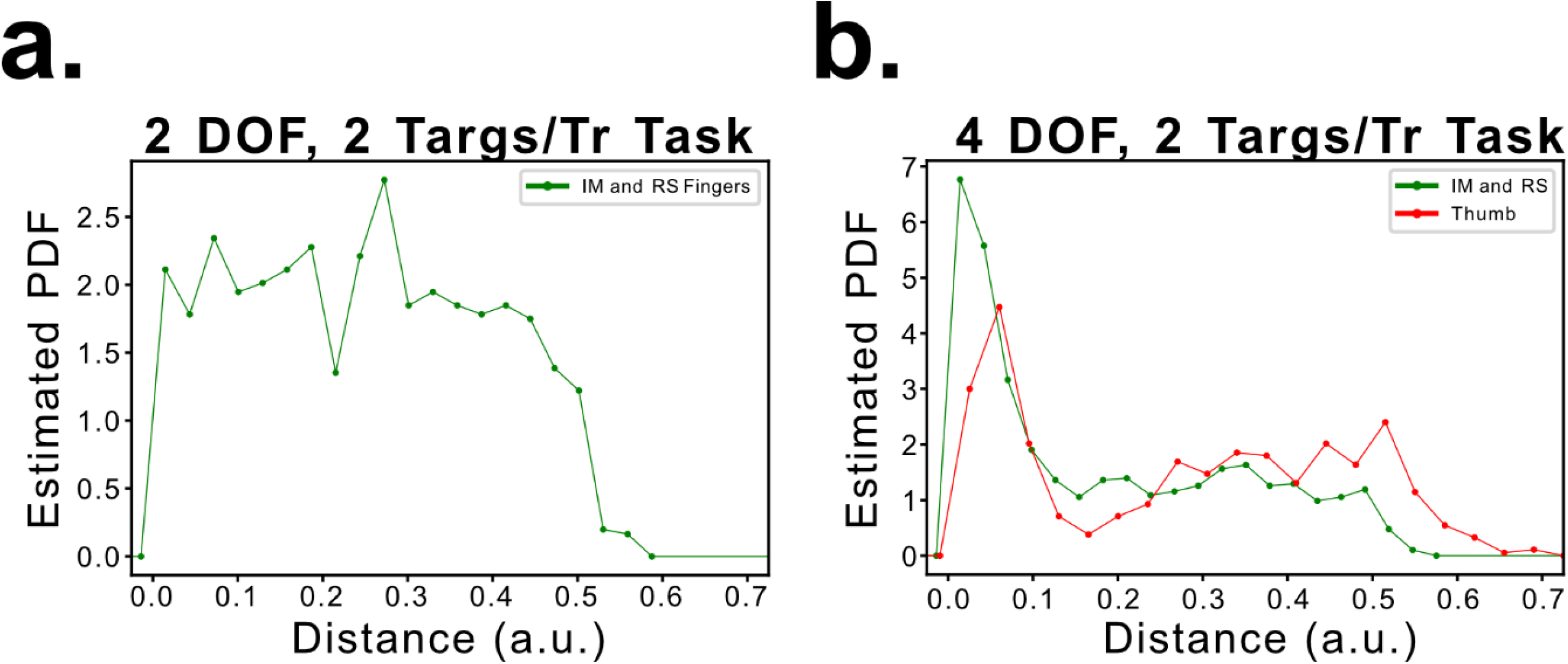
Distribution of finger distances. For the 2D and 4D task with 2 new targets/trial (in Figs. 2b, 2d), the distribution of distances for successful trials for the **(a)** 2D decoder and **(b)** 4D decoder. The histogram is normalized so that the area under the curve equals 1. In **a**, the green curve represents the distance for index-middle (IM) finger and ring-small (RS) finger flexion/extension combined over all trials for the 2D decoder and task trials in Fig. 1e. In **b**, the green curve represents combined IM finger and RS finger flexion/extension distances, and the orange curve represents the 2D distance for the thumb (combining the 2D components of flexion/extension and abduction/adduction) for trials using the 4D decoder and task in Fig. 1e. a.u., arbitrary units; DOF, degree of freedom; PDF, probability distribution function.

**Extended Data Fig. 5.**
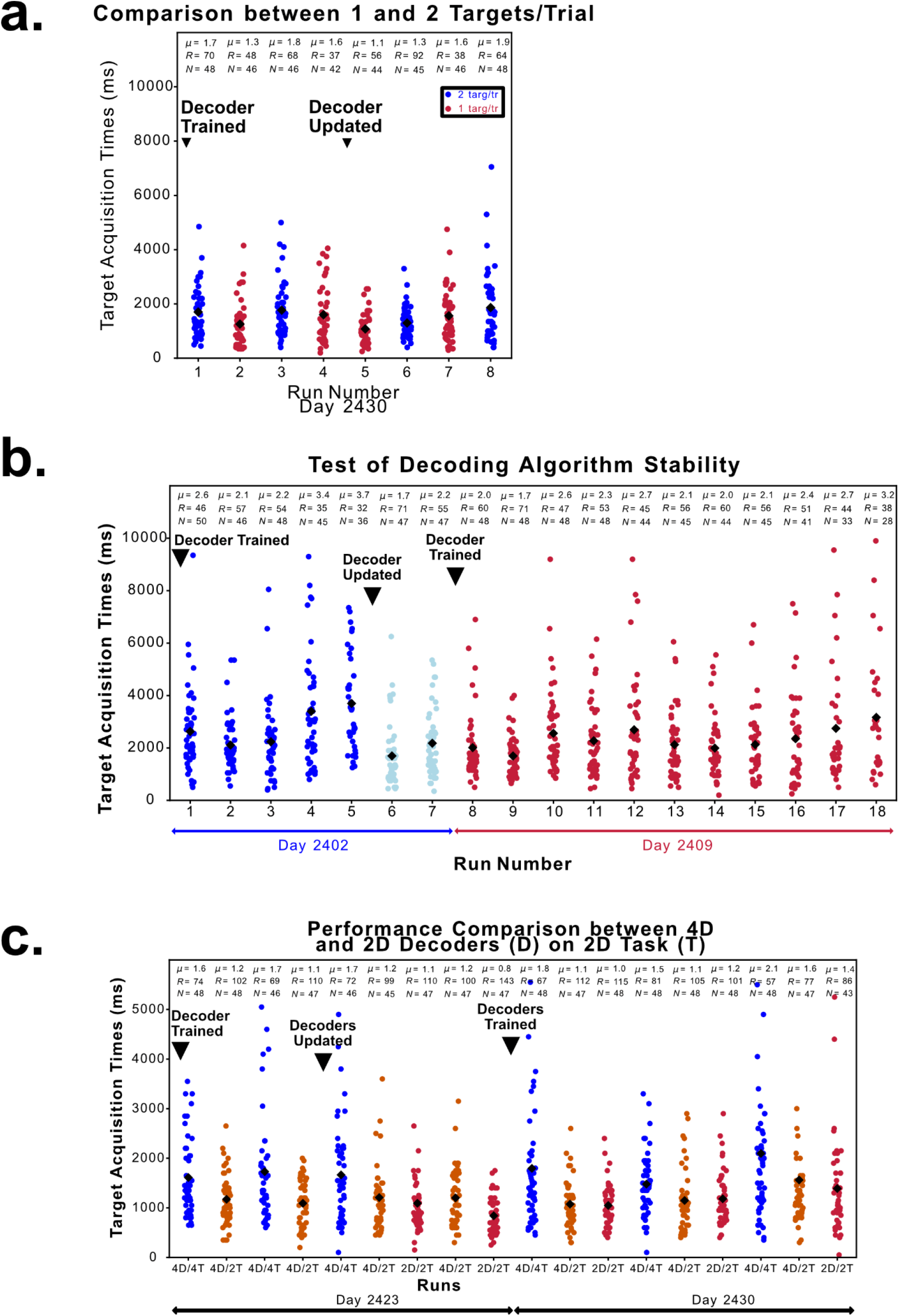
Target acquisition times for a variety of comparisons. **a**, Performance comparison for 1 vs. 2 new targets/trial (targ/tr). Population data for the target acquisition times (ms) using the 4D decoder for 2 new targets/trial (blue) and 1 new target/trial (red) over blocks of the task. Each dot corresponds to a trial, and the black diamond indicates the mean value. *N* denotes the number of trials per block, *μ* denotes the mean value, and R denotes the rate of targets acquired in targets/min. **b**, Decoder stability test for 2 trial days using the 4-DOF decoder for 2 new targets/trial. For days 2402 (blue and light blue) and 2409 (red), the decoder was trained (upside down triangle with “Decoder Trained”) and then used in consecutive blocks until trials could not be reliably completed. On day 2402, the decoder was re-trained (“Decoder Updated”) to demonstrate the recovery of performance on 2 subsequent blocks (light blue). **c**, The 4D and 2D decoders on 2D task. Target acquisition times (ms) for the 4D and 2D decoders are compared on the 2D task with 2 new targets/trial task (2T). The 4D decoder is also run on the 3-finger group, 4D task (4T) with 2 new targets/trial. The blocks on the 2423 and 2430 trial days of data collection represent consecutive blocks without re-training unless otherwise indicated. The labels 4D/4T indicate the 4D decoder run on 4T; 4D/2T indicates the 4D decoder on 2T; and 2D/2T indicates the 2D decoder on 2T.

**Extended Data Movie 1**: The 2D decoder on the 2D task with a mean acquisition time of 0.84 ± 0.05 s, corresponding to a target acquisition rate of 142 targets/min.

**Extended Data Movie 2**: The 4D decoder on the 4D task with 2 new targets/trial with a mean acquisition time of 1.30 ± 0.08 s, corresponding to a target acquisition rate of 92 targets/min.

**Extended Data Movie 3**: The 4D decoder on the 4D task with 1 new target/trial with a mean acquisition time of 1.08 ± 0.09 s, corresponding to a target acquisition rate of 56 targets/min.

**Extended Data Movie 4**: The 4D decoder on the 2D task when 2 DOF are fixed and not allowed to move (thumb abduction/adduction and ring-small flexion/extension). The mean acquisition time was 1.07 ± 0.06 s, corresponding to a target acquisition rate of 112 targets/min.

**Extended Data Movie 5**: Exemplar block of the finger intracortical brain-computer interface translated to control a quadcopter with 4 DOF during an obstacle course presented in Fig. 4b and flight path shown in Fig. 4c.

**Extended Data Movie 6**: Using the finger intracortical brain-computer interface translated to quadcopter control to navigate through randomly appearing rings to demonstrate spontaneous, free-form control.

